# Mechanism for evolution of diverse autologous antibodies upon broadly neutralizing antibody therapy of people with HIV

**DOI:** 10.1101/2025.03.05.641732

**Authors:** Deepti Kannan, Eric Wang, Steven G. Deeks, Sharon R. Lewin, Arup K. Chakraborty

## Abstract

Antiretroviral therapy (ART) inhibits Human Immunodeficiency Virus (HIV) replication to maintain undetectable viral loads in people living with HIV, but does not result in a cure. Due to the significant challenges of lifelong ART for many, there is strong interest in therapeutic strategies that result in cure. Recent clinical trials have shown that administration of broadly neutralizing antibodies (bnAbs) when there is some viremia can lead to ART-free viral control in some people; however, the underlying mechanisms are unclear. Our computational modeling shows that bnAbs administered in the presence of some viremia promote the evolution of autologous antibodies (aAbs) that target diverse epitopes of HIV spike proteins. This “net” of polyclonal aAbs could confer control since evasion of this response would require developing mutations in multiple epitopes. Our results provide a common mechanistic framework underlying recent clinical observations upon bnAb/ART therapy, and they should also motivate and inform new trials.

## INTRODUCTION

HIV is a highly mutable virus that is not controlled by the immune responses of most people living with HIV, and for which there is no effective vaccine.^1^ Highly effective ART can completely suppress HIV replication in most people.^2^ However, ART does not provide a cure, since integrated viral DNA encoding replication-competent virus persists indefinitely within transcriptionally silent infected cells (the “latent reservoir”).^3^ Discontinuing ART results in viral rebound within two to three weeks in nearly all people with HIV. ART therefore needs to be administered for life, which poses significant challenges.^4^ There is thus a global interest in developing strategies that result in either full eradication of the virus or sustained ART-free viral control.^5–7^

One such therapeutic strategy that has been tested in clinical trials is the passive administration of broadly neutralizing antibodies (bnAbs).^8–15^ Potent bnAbs have been isolated from some people with HIV.^16–20^ They neutralize diverse strains of HIV-1 because they bind to regions (epitopes) on the spike envelope (Env) trimer that are relatively conserved across strains; for example, the CD4 binding site which binds to the CD4 human receptor to propagate infection.^21^

Recent clinical trials suggest that bnAb administration in the presence of some viremia can induce ART-free viral control in some participants.^9,11^ Importantly, control persists long after levels of bnAbs decline to sub-therapeutic levels. For example, in the eCLEAR trial, 3BNC117 (a CD4 binding site bnAb) and ART were administered at roughly the same time in participants newly diagnosed with HIV infection. After antiretroviral treatment interruption (ATI), 4 of the 5 participants who harbored 3BNC117-sensitive virus maintained ART-free viral control.^11^ Similarly, in the TITAN trial, where 3BNC117 was administered with another bnAb (10-1074) at the time of ATI, 4 of 11 participants met the criteria of ART-free viral control.^9^ In both these trials, participants had some level of viremia at the time bnAbs were administered. In contrast, in the ROADMAP trial, where 3BNC117 was administered during stable ART treatment (undetectable viremia), viral rebound occurred rapidly post-ATI in all participants.^10^ Understanding the mechanisms that conferred post-ATI control in some participants when bnAbs were administered with ongoing viremia is important for developing an effective cure intervention.

The leading proposed mechanism for ART-free viral control post-bnAb therapy is a “vaccinal” effect: bnAbs bind to viral proteins in circulation to form immune complexes, which when ingested by dendritic cells (DCs), prime them to more effectively present antigen^22–24^ and activate virus-specific CD8+ T cells, as evidenced by data from non-human primate models.^25,26^ In humans, enhanced HIV-specific CD8+ T cell responses were observed in early proof-of concept studies^27^ and a modest increase was seen in the eCLEAR trial.^11,28^ However, there was no evidence of a vaccinal effect in the TITAN study.^9^ CD8+ T cell responses are more effective for those with protective HLA alleles (e.g., HLA-B27, HLA-B57, HLA-B*58).^29–31^ However, many ART-free viral controllers post ATI, including 3 of the 4 in the eCLEAR trial,^11^ did not carry protective HLA alleles.^32^ Thus, other mechanisms besides CD8+ T cell responses may contribute to ART-free viral control.

Upon infection, autologous antibodies (aAbs) continually develop in response to circulating virus, and HIV evades these responses by evolving low fitness cost escape mutations in the targeted Env epitopes.^33– 35^ However, aAbs have recently been recognized as potentially important for HIV control.^36–38^ Several studies have shown that aAbs can evolve during ART,^36,37^ and that these responses can contribute to ART-free viral control, even in the absence of an additional intervention.^36,39,40^ Potent, non-neutralizing antibody effector functions such as antibody-dependent cellular cytotoxicity (ADCC) can also develop during ART^41^ and may be important for ART-free viral control.^42,43^ In particular, the VISCONTI study showed that transient episodes of viremia correlated with strong cross-neutralizing antibody responses and non-neutralizing effector functions in post-treatment controllers (PTCs).^44,45^ This study bears resemblance to the eCLEAR and TITAN trials, where viremia at the time of therapy is linked to ART-free viral control post therapy.

New clinical data also suggest that aAbs could be relevant for ART-free viral control post bnAb therapy. Preliminary analysis of data from the BEAT2 trial suggests that aAb responses upon combination bnAb therapy contributed to ART-free viral control in a subset of participants.^46^ Importantly, Schoofs et al. showed that bnAb therapy enhanced host neutralizing antibody responses, as evidenced by increased neutralization of both autologous and heterologous viruses 6 months after therapy.^14^

Several key questions emerge. Could bnAbs promote the evolution of potent aAb responses, and if so, how? Why is some accompanying viremia at the time that bnAbs are administered important for the evolution of such a response? How might aAbs acquire sufficient breadth to contribute to ART-free viral control, despite prevailing virus diversity?

We address these questions by adapting past computational models of the humoral immune response. Our results reveal the following mechanism: (1) Potent bnAbs bind to and deposit diverse circulating virions on Follicular Dendritic Cells (FDCs), thus exposing diverse spike protein epitopes that can drive Germinal Center (GC) reactions. (2) The first wave of immunodominant antibodies that evolve can subsequently mask their epitopes,^49,54–56^ which along with the presence of sufficient amounts of antigen on FDCs, promotes the evolution of new subdominant antibody responses that target other epitopes. Targeted epitopes are successively masked and antibodies targeting diverse epitopes develop during bnAb therapy/ART while viral replication and mutation is low. The evolution of such a polyclonal “net” of antibodies could delay viral rebound once ART is stopped since escape mutations would have to develop in multiple epitopes in order for the prevailing viruses to evade such a response. Our results show that only an intermediate level of viremia at bnAb therapy initiation promotes the evolution of a polyclonal net of antibodies. This finding may explain why such responses may have developed in participants with ART-free viral control in the eCLEAR and TITAN trials, as well as those who had viremic episodes in the VISCONTI study, but not in the participants of the ROADMAP trial.

We hope that the mechanism that we propose for the evolution of aAbs upon bnAb therapy, and their role in conferring ART-free viral control, will be interrogated further in clinical trials and inform cure strategies.

## RESULTS

### Model of Germinal Center reactions and generation of antibodies

The computational method we develop to simulate the humoral immune response to bnAb therapy is adapted from a model we employed to study antibody responses upon sequential immunization with SARS-CoV-2 vaccines.^49^ Below we summarize the main biological steps that we simulate (Fig. 1) and outline how the steps are implemented in the computational model (see STAR Methods for details).

**Figure 1:**
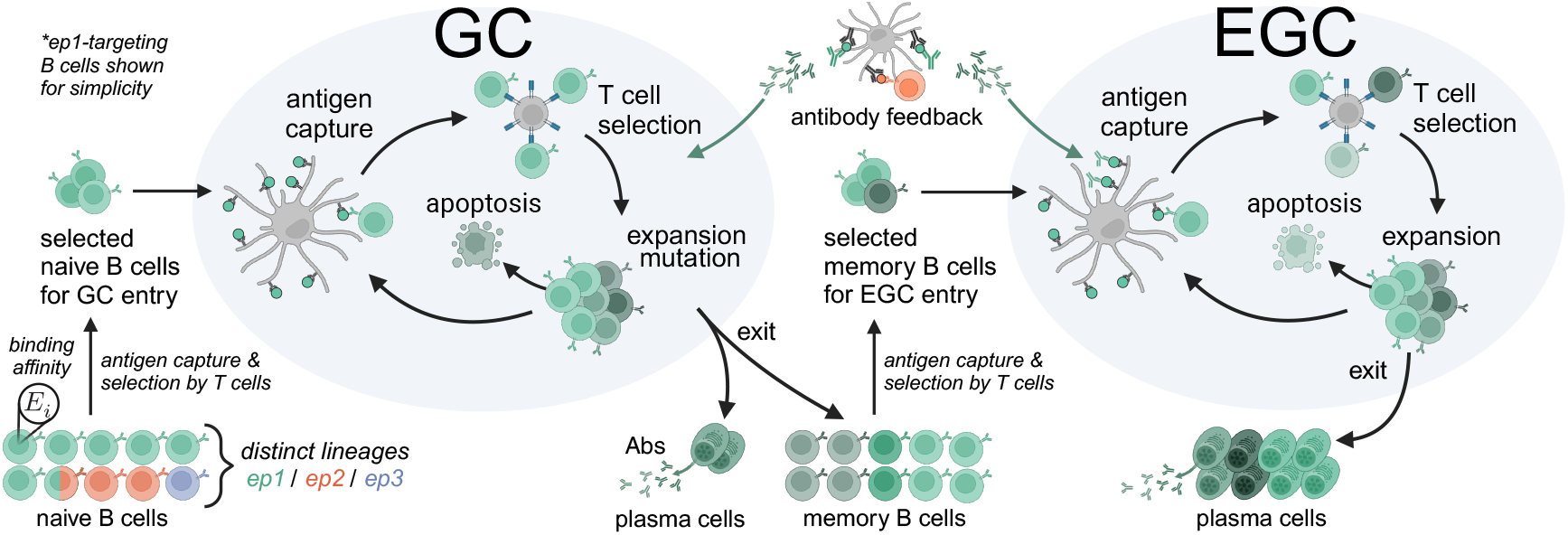
Biological steps simulated in computational model of the humoral immune response. Naive B cells targeting different epitopes are initialized with binding affinities, *E*_*i*_. Naive B cells are activated by antigen capture and positively selected by T cells for GC entry. GC B cells undergo rounds of selection, proliferation, mutation, and apoptosis. Nonlethal, nonsilent mutations result in a change in binding affinity. Mutant clones are depicted with different shades of green. A small fraction of GC B cells are exported as antibody-secreting plasma cells and memory B cells. Memory B cells are selected for entry into compartments outside GCs (referred to as EGC) where they undergo affinity-dependent expansion in a similar manner as in the GC but without mutation. The EGC exports plasma cells, which produce most of the antibodies in serum. Antibodies perform feedback regulation of the humoral response by entering ongoing GCs/EGCs and masking their target epitopes on FDCs. The model predicts the dynamics of the different B cell types, and the concentration and affinity of antibodies.

Since bnAbs are administered in relatively high concentrations and have a high affinity to diverse circulating strains that are bnAb sensitive, we assume bnAbs quickly form immune complexes with these strains and deposit them on FDCs. Since antigen is retained on FDCs for long times,^57,58^ we further assume a fixed total antigen concentration ([Ag]) throughout the course of the simulation. Long-term antigen availability implies a persistent GC response.^59^ We also assume that [Ag] on FDCs is high enough that GC B cells can potentially interact with all epitopes in the diverse antigens displayed on FDCs.

In our model, each epitope on the HIV spike proteins that is displayed on FDCs is associated with a distinct population of germline/naive B cells that can target it. Epitopes that are shared across multiple variant strains are targeted by the same germline population. Naive B cells with germline affinity above a threshold of *K*_*d*_ = 10^−6^*M* can be activated by antigen encounter.^60–62^ The affinity distributions of germlines targeting different epitopes are defined such that more immunodominant epitopes have higher affinity tails than less dominant epitopes. We also vary the fraction of naive precursor lineages that target each epitope, with more precursors for more immunodominant epitopes.

We then employ agent-based simulations of GC reactions, affinity-dependent expansion of memory B cells outside GCs, and production of antibodies for 400 days of treatment (as in recent clinical trials^9,11^). At each simulation time step, naive B cells are activated with a probability dependent on the amount of antigen they internalize,^63^

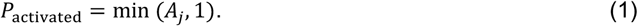

The amount of antigen, *A*_*j*_, that the *j*^*th*^ B cell internalizes increases with its binding “affinity”,^61,64^ defined as *E*_*j*_ = −log^10^ *K*_*d*_, and then saturates at the affinity ceiling of *E*_*j*_ = 10.^60^ The amount of internalized antigen, *A*_*j*_, also increases with the concentration of the B cell’s target epitope *i* on FDC, *C*_*i*_,

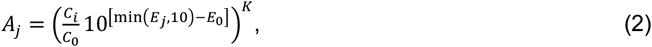

where *E*_0_ = 6 corresponds to the minimum affinity for activation.^61,62^ The reference concentration *C*_0_ controls how sensitive *A*_*j*_ is to changes in epitope availability on FDCs. Higher values of *K* imply more stringent selection with respect to differences in binding affinity.

Activated B cells then compete for positive selection signals from T follicular helper cells (Tfh) in order to enter GCs.^65,66^ The probability of being positively selected depends on the relative amount of antigen internalized by a B cell compared to the average amount internalized by other activated B cells, and the relative number of Tfh cells, *N*_Tfh_, compared to the total number of activated B cells, *N*_Bcells_. The rate of positive selection for a B cell, *j*, is given by

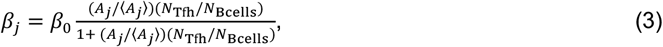

where the parameters *N*_Tfh_ and *β*_0_ are chosen to control the rate of GC entry.

Positively selected B cells continuously enter GCs,^64,67^ where they undergo rounds of selection, proliferation, and mutation. Selection of GC B cells occurs in two steps, antigen internalization (Equations 1-2) and competition for T cell help (similar to Equation 3).^68^ The maximum rate of positive selection from T cell help for GC B cells is set to match the observation that GC B cells typically divide a maximum of 4 times in a day.^69^ In addition, the number of Tfh cells in the GC is chosen based on measurements from fine needle biopsies of people with HIV^70^ and assumed to be constant during therapy. At each time step, positively selected GC B cells duplicate, and one of the daughter cells is re-cycled for further rounds of selection. Of the other daughter cells, 5% exit the GC as antibody-secreting plasma cells or memory B cells^71^ and the rest can mutate (50% of mutations are silent, 30% are lethal, and 20% are affinity-changing^72^). The change in a GC B cell’s binding affinity, Δ*E*_*j*_, is drawn from a log normal distribution, with a 5% chance of increasing the affinity and a 95% chance of decreasing the affinity (unpublished work, Victora Lab).^72^ Our model reproduces the experimental observation that B cells that receive stronger selection signals from Tfh cells (larger *β*_*j*_) will divide a larger number of times.^73^ All GC B cells stochastically undergo apoptosis at a constant rate.^74^ This cycle of selection, export, mutation, and death repeats in every time step.

For each parameter set, we simulate 200 GCs, based on the observation of 100 GCs in 6 micrometer sections of lymphoid tissues in mice.^75^ The memory B cells generated by all GC reactions are pooled together and are continuously activated outside GCs according to the same activation probability as that of naive B cells (Equation 1) and then compete for T cell help according to Equation 3. The selected memory cells undergo antigen affinity-dependent selection and expansion in compartments outside GCs that we refer to as “Extra Germinal Centers” (EGCs).^53,76,77^ The same processes occur in the EGC as in GCs, but with little to no mutation because these cells express very low levels of AID.^78^ Consistent with experimental data,^76^ 60% of EGC memory B cells differentiate into plasma cells which secrete antibodies. The concentration of aAbs at each time step is determined by the number of plasma cells in circulation and the rate of antibody secretion. The affinity of these antibodies is taken to be the mean affinity of the plasma cells from which aAbs are generated.

Several mouse studies have shown that epitope masking restricts new B cell responses to the same epitope while increasing responses to different epitopes,^54,56,79^ and this phenomenon has also been observed in human clinical data.^55,80^ Thus, aAbs generated by the GC/EGC response in our simulations mediate feedback regulation of the subsequent GC/EGC processes via epitope masking. That is, antibodies can bind to their epitopes *i* on FDCs and partially mask them, reducing *C*_*i*_ and increasing the relative concentration of subdominant epitopes. Because viral replication is low during ART and bnAb therapy, we neglect potential coevolution of the virus as well as changes to antigen presentation on FDCs due to newly generated aAbs.^49,50^

### BnAb-mediated presentation of diverse circulating strains on FDCs and epitope masking promote sequential evolution of antibodies targeting different epitopes

If there is some viremia when bnAbs are administered, these potent and cross-reactive antibodies can bind to and deposit diverse circulating strains on FDCs. We employ our stochastic simulations to study the humoral response to the diverse epitopes across multiple antigens that are thus accessible to B cells during the treatment phase. For ease of illustration of concepts, we first consider a simple case where three non-overlapping epitopes that are differentially immunodominant are displayed on FDCs at equal concentrations.

Figures 2A and 2B show that at early times only the most immunodominant response emerges, as quantified by the rise in the concentration and affinity of antibodies targeting epitope 1 (green). As time ensues, our results show that the concentration and affinities of antibodies targeting the subdominant epitopes also increase, and a polyclonal net of antibodies targeting different epitopes emerges. Our study reveals the following mechanism underlying these results.

**Figure 2:**
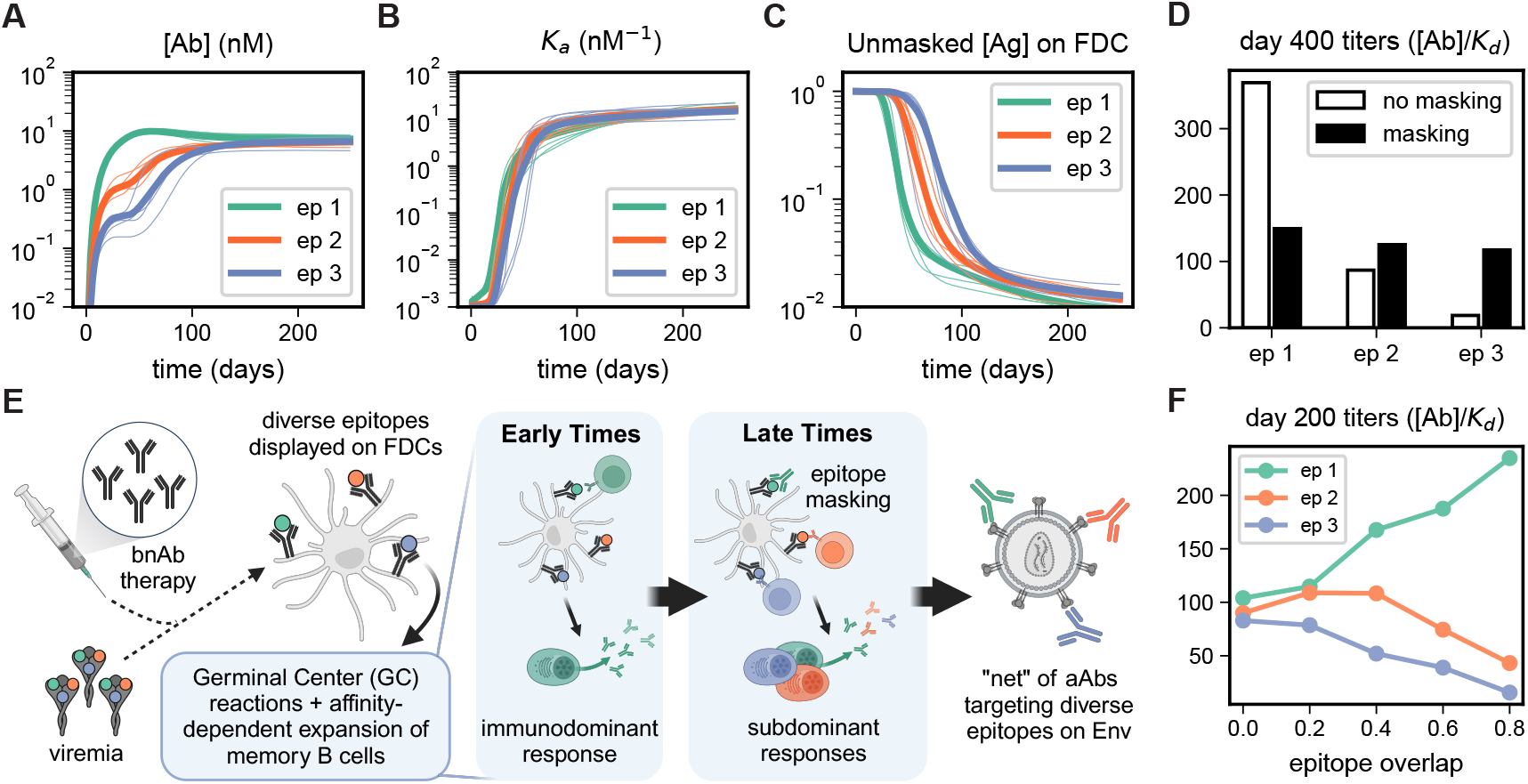
Broadly neutralizing antibody (bnAb) therapy promotes sequential evolution of autologous antibodies (aAbs) targeting different epitopes during treatment. Simulated dynamics of concentrations (A) and affinities (B) of antibodies targeting 3 distinct epitopes (ep1, ep2, ep3) arranged in an immunodominance hierarchy. Thick lines show the mean over 50 individual simulations (select 5 shown in thin lines). (C) Newly generated antibodies bind to their target epitope on FDCs and reduce its effective concentration. Masking of each epitope proceeds sequentially according to the immunodominance hierarchy. (D)Ablating epitope masking in our simulations increases titers of aAbs targeting the immunodominant epitope and decreases responses to subdominant epitopes. (E)According to our proposed mechanism, bnAbs bind to diverse circulating strains and deposit them on FDCs, leading to a sustained GC response. At early times, the most immunodominant response dominates. These antibodies then mask epitope 1 on FDC, promoting the evolution of subdominant responses. Over time, antibodies targeting different epitopes can evolve sequentially during the treatment phase. (F)Antibody titers in the case where epitope 1 overlaps with epitopes 2 and 3, meaning antibodies targeting epitope 1 will also partially mask the other two epitopes. As the level of epitope overlap increases, the titers of antibodies targeting epitopes 2 and 3 decrease, while titers to the dominant epitope increase.

The initial antibodies targeting the immunodominant epitope bind to epitope 1 on FDCs (epitope masking), thus reducing its effective concentration and increasing the relative concentration of epitopes 2 and 3 (Figure 2C). As a consequence of this shift in the balance of epitope availability on FDCs, more naïve B cells targeting subdominant epitopes enter GCs (Figure S2A) and compete effectively. As a result, more GC B cells targeting subdominant epitopes are positively selected and exported as memory and plasma cells (Figure S2B-D). High affinity subdominant memory cells are then expanded outside GCs, increasing the number of plasma cells targeting subdominant epitopes and reducing the number of plasma cells that target epitope 1 (Figure S2E-F). As a result, production of antibodies targeting epitope 1 slows down slightly at around day 50, while the concentration and affinity of antibodies targeting epitopes 2 and 3 increase (Figure 2A). Eventually, the concentrations of all 3 antibody populations reach a steady state determined by a balance of plasma cell differentiation and death rates, as do the affinity distributions of memory B cells (Figure S1B-C). The mean affinity of all 3 antibodies plateaus at the imposed affinity ceiling of 10^1^nM^−1^ (Figure 2B).^60^

The potency of the serum antibody response can be quantified by the binding titer, which is the product of the antibody concentration and its affinity (1*/K*_*d*_). When we turn off the effect of epitope masking in our simulations (Figure 2D), we find that the antibody response is dominated by the most immunodominant response, and balanced titers of antibodies targeting different epitopes do not result. Thus, epitope masking is the key mechanism driving the sequential evolution of a potent antibody response targeting diverse epitopes deposited by bnAbs on FDCs (Figure 2E).

Such a net of antibodies targeting diverse epitopes would be highly unlikely to develop during natural infection. Since viral load and replication are high in the absence of ART, mutations in HIV spike proteins that incur relatively low fitness cost would evolve to evade the initial immunodominant antibody response before subdominant antibody responses emerge.

Each of the thin lines in Figures 2A-C corresponds to a simulation of 200 GCs and one EGC, representative of an *in silico* “person”. The stochastic variation in the timing and strength of the antibody response across simulations could explain some of the variation in the time to rebound following ART cessation among participants in clinical trials. In our simulations, we assume that the antigen concentration on FDCs remains constant during ART. Differences in the duration of antigen retention on FDCs would also lead to variations in the number of aAb responses that develop. Another factor that could influence the timing and strength of the antibody response is the degree of spatial overlap between epitopes on the HIV spike proteins. If there is epitope overlap, antibodies targeting the most dominant epitope will also partially mask overlapping subdominant epitopes. As epitope overlap increases, titers to subdominant epitopes are suppressed at day 200 (Figure 2F). Epitope overlap can delay or abrogate the evolution of subdominant responses.

Our proposed mechanism can be tested by mapping the epitopes of aAbs generated in participants who achieve ART-free viral control following bnAb therapy. Our results suggest that participants who achieved control and initially harbored a diversity of circulating strains should exhibit multiple aAb responses directed toward epitopes that are more likely to be non-overlapping.

### An optimal range of viremia when bnAbs are administered promotes development of the net of antibodies targeting diverse epitopes

The concentration of antigen displayed on FDCs depends on the viral load at the time of bnAb administration, the concentration of bnAbs, the affinity of bnAbs for the circulating strains, and the kinetics of transport to FDCs.^49,81^ For a fixed concentration and affinity of bnAbs, the antigen concentration on FDCs is directly proportional to the viral load at the time of therapy initiation.

Again, consider the case of three distinct epitopes that are differentially immunodominant (Figure 3). When viremia is low when bnAbs are administered, less antigen will be displayed on FDCs. At low antigen concentrations, no potent antibody responses develop, since there is insufficient antigen to enable naive B cells to enter GCs. At somewhat larger antigen concentrations, selection is more stringent and so significant antibody titers develop only for the immunodominant response. This result may explain why viral rebound occurred rapidly in the ROADMAP trial,^10^ where bnAbs were administered under stable ART with undetectable viral load. New aAbs would either have not developed at all, or those targeting a few immunodominant epitopes would have developed and then been evaded by mutation upon ATI.

**Figure 3:**
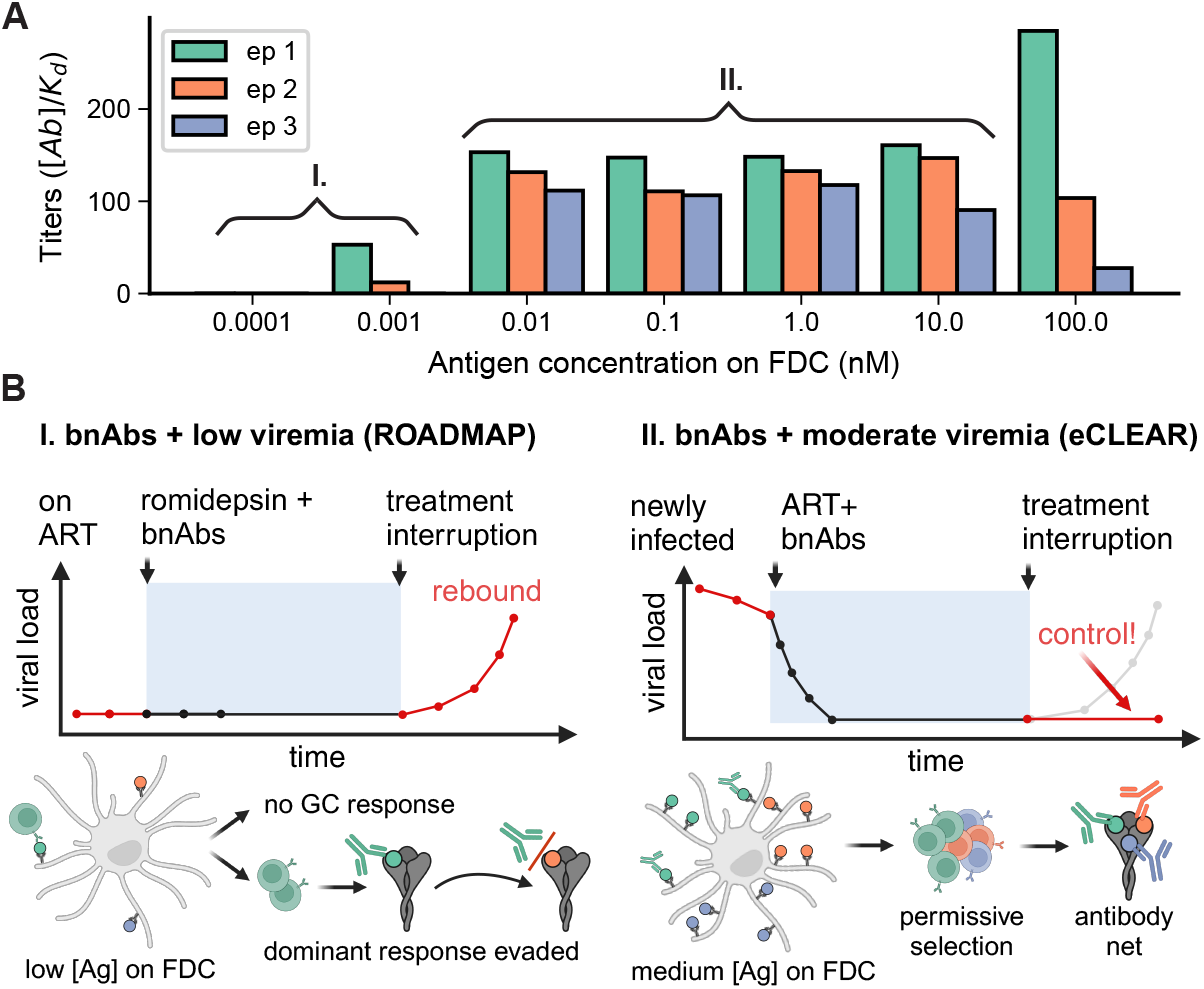
An optimal range of antigen concentration on FDCs promotes balanced antibody titers to diverse epitopes. (A)Titers of aAb responses to three epitopes as a function of total concentration of antigen on FDCs, [Ag]. (B)If [Ag] is low on FDCs, either no GC response develops or only the most immunodominant response develops. This could be the case in the ROADMAP trial^10^, where bnAbs were given to people on stable ART. If [Ag] is in an intermediate range, more subdominant responses can evolve, as may be the case in for example the eCLEAR trial^11^, where bnAbs were given to people with detectable viremia. Only immunodominant responses develop when [Ag] is in excess of the newly generated aAbs because of ineffective epitope masking.

When antigen concentration on FDCs is in excess of the generated antibodies, the immunodominant response dominates since aAbs cannot mask a significant fraction of the immunodominant epitope to enable the development of subdominant responses. Only when the amount of antigen on FDCs is in an intermediate range do we find that balanced titers to diverse epitopes develop (Figure 3). This result holds as we vary the unknown model parameter, *C*_0_ (Figure S3).

Our results could explain why some participants achieved ART-free viral control in the eCLEAR^11^ and TITAN^9^ trials, where participants had some viremia and thus moderate levels of antigen on FDCs at the time of bnAb administration. Our results may also explain why only the PTCs that experienced low-level viremic episodes exhibited strong cross-neutralizing antibody responses in the VISCONTI study.^45^ Antibodies that developed during a viremic episode could display moderate levels of antigen on FDCs, leading to the development of more aAbs. These antibodies could then mask their corresponding epitopes, promoting the evolution of other aAbs that target subdominant epitopes. This effect would be amplified upon subsequent viremic episodes.

The development of new aAb responses upon bnAb therapy has been observed clinically by Schoofs et al.^14^ Measurements of the IgG response 24 weeks after 3BNC117 bnAb administration showed that neutralization of both pre-treatment virus and autologous week 24 virus increased compared to the pre-treatment IgG response. This effect was was more pronounced for participants who were off ART at the time bnAbs were administered, which is consistent with our prediction that some viremia promotes evolution of diverse aAbs. In addition, sera from participants in the trial exhibited significant increases in neutralization to multiple heterologous viruses. The breadth of the response suggests that antibodies targeting multiple, distinct epitopes may have evolved, as predicted by our *in silico* results.

### Viral diversity, Tfh counts, and B cell germline characteristics influence the potency and size of the antibody net

Next we studied how the diversity of circulating strains prior to bnAb therapy affect the aAb response. Consider a simple case where there are two variant antigens with six epitopes each for a total of 12 possibly distinct epitopes (if the strains are maximally divergent). In Fig. 4A, we vary the number of shared epitopes across the two antigens, which is a metric of the diversity of circulating strains pre-bnAb therapy. We assume that the shared epitopes are less immunodominant than variable epitopes.

**Figure 4:**
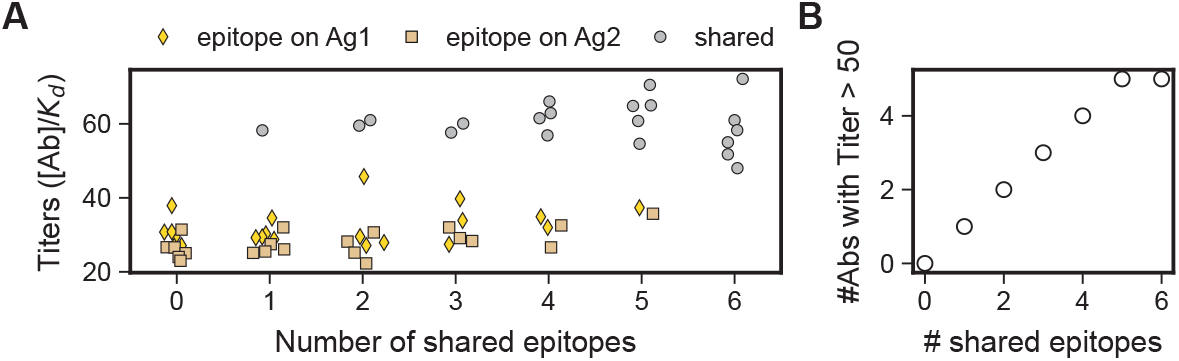
Diversity of circulating strains at the time of bnAb therapy affects the potency and number of aAb responses. (A)Titers of antibodies that develop towards two antigens with six epitopes each. The abcissa represents the number of shared epitopes between the 2 antigens. Epitopes that are unique to antigen 1 (yellow diamond) or antigen 2 (tan square) are assumed to be more immunodominant than shared epitopes (grey circle). Titers to shared epitopes are higher than those targeting unique epitopes despite being less immunodominant. (B)As the number of shared epitopes increases, the number of potent aAb responses increases.

In Figure 4A, each graphed point represents the antibody titer directed toward a particular epitope present in the antigen cocktail. If both antigens are present in equal concentrations, then shared epitopes are present on FDCs at double the abundance of the variable ones. As a consequence, we find that antibody titers directed toward shared epitopes are higher than to epitopes that are not shared among variants, despite shared epitopes being less immunodominant (Figure 4A). Indeed, the number of distinct epitopes present on all strains that are potently targeted increases with the number of shared epitopes (Figure 4B). Our results imply that a lower diversity of circulating strains present at the time of bnAb/ART therapy initiation should lead to potent aAb responses targeting a larger number of shared epitopes on diverse strains. A larger “net” of such aAbs will be harder for the virus to escape upon ATI, since mutations in a larger number of shared epitopes would have to evolve. Thus, PTC via aAbs is more likely in people who start ART early after infection, such that circulating strains and those that emerge from the latent reservoir post-ATI are not too divergent. This prediction is consistent with clinical observations showing that PTC correlates with stronger antibody responses and lower provirus diversity in the latent reservoir.^36^

Another immune parameter affecting the aAb response is the number of activated Tfhs. According to the “vaccinal” effect hypothesis, DCs are more effectively primed by bnAb-antigen immune complexes,^22,23^ resulting in enhanced activation of T cells, including Tfh cells (Figure S4A).^24^ In our simulations, we find that increasing the number of Tfh cells within each GC increases the binding titers to all three epitopes (Figure S4B). This is because more T cell help implies more positively selected B cells and larger GC sizes (Figure S4C).^70^ Similarly, increasing Tfh in the EGC increases the overall scale of the antibody titers and leads to more balanced titers across epitopes (Figure S4D-G). This is because memory cells compete for productive interactions with a larger number of T cells, leading to more permissive selection of subdominant responses. Thus, our model predicts that individuals with higher Tfh counts will be more likely to evolve potent aAbs that can confer ART-free viral control.

Characteristics of the germline B cell populations also influence the timing of when antibody responses evolve. For example, increasing the gap in affinity between precursors targeting epitope 2 and epitope 3 delays the emergence of significant titers towards epitope 3 (Figure S5A). This is because it takes longer for naive B cells targeting the subdominant epitope to get activated, enter GCs, and acquire sufficient mutations to outcompete the immunodominant GC B cells. Similarly, more balanced frequencies of precursors targeting different epitopes shortens the delay time between aAb responses (Figure S5B). These variations in germline B cell characteristics could explain some of the variation in outcomes between people on bnAb therapy.

### Clonal dynamics of B cells provide insights into evolution of aAbs targeting different epitopes

We next examined the clonal dynamics of B cells within and outside GCs in order to further explore the mechanism by which a polyclonal net of aAbs can evolve. For the simulations with three epitopes, there are 2000 Env-specific naive B cell lineages, where 65% target immunodominant epitope 1, 25% target epitope 2, and 10% target epitope 3. We track which clones enter GCs, and monitor how the frequencies of each lineage within a GC evolve over time.

In Figure 5A, we illustrate the clonal dynamics in three representative GCs. Within the GC, there is both interclonal competition within an epitope class, and interclass competition between populations targeting different epitopes. At early times, the GC has a multi-clonal structure with most clones targeting epitope 1. Over time, due to sequential masking of epitopes, more clones targeting subdominant epitopes can enter the GC. Still, in 66% of GCs, exemplified by the first row of Figure 5A, a single clone targeting the immunodominant epitope takes over the GC, reaching a frequency of nearly 80% (Figure 5B, green line). In 23% of GCs (second row, Figure 5A), a subdominant clone targeting epitope 2 takes over the GC, and in 11% of GCs (third row, Figure 5A), a clone targeting epitope 3 dominates the GC. The mean time at which the winning clone takes over the population is delayed for subdominant epitopes (Figure 5B), but is highly stochastic.

**Figure 5:**
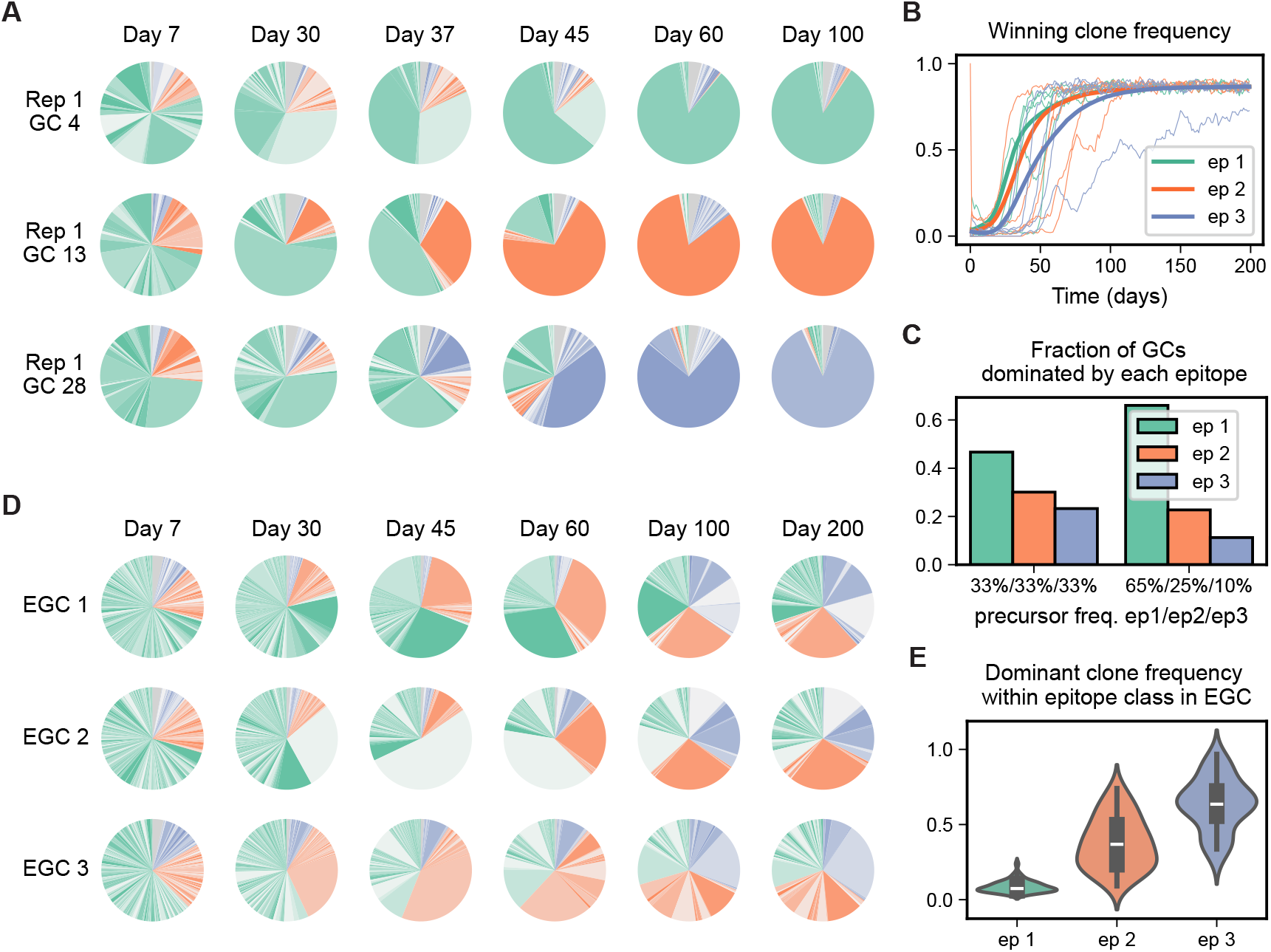
Clonal dynamics of B cells within and outside GCs reveal how polyclonal antibody response emerges. (A)Each pie slice represents a different clonal lineage within a GC, where shades of green correspond to clones targeting epitope 1, orange for epitope 2 and purple for epitope 3. Over time GCs become more monoclonal, with the most dominant clone targeting epitope 1 most of the time (top row), but sometimes targeting epitope 2 (second row) or 3 (third row). (B)Frequency of the clone that dominates the GC at day 400 as a function of time. Thick lines represent averages over the subset of GCs where a particular epitope population dominates. Thin lines show trajectories from 5 GCs in each epitope class. (C)Fraction of GCs where the population targeting each epitope wins the evolutionary competition. Distribution is more balanced when the precursor frequencies of naive B cells targeting each epitope are equal (left cluster of bars) as compared to when they are unequal (right cluster of bars). (D)Clonal dynamics of memory B cells in three representative EGCs. (E)Distribution of frequencies of the most dominant clone within each epitope-targeting population at day 400 across 50 EGCs. The immunodominant population shows the most clonal diversity.

The fittest clone eventually taking over the population is expected since the GCs can avoid stochastic extinction due to sustained antigen and Tfh cell availability. In reality, antigen on FDCs is depleted over time as it is internalized by B cells and due to other decay processes. Therefore, takeover of any GC by a single population of B cells is unlikely. The more likely situation is reflected by the results in Figure 5A-B at time points earlier than day 100.

The fraction of GCs in which a particular population wins is determined by the germline immunodominance hierarchy. For example, if the precursor frequencies of naive cells targeting the 3 epitopes are equal, the distribution of GCs targeting each epitope is more balanced, with the differences arising from different germline affinity distributions for the three epitopes (Figure 5C).

The results noted above explain why the plasma and memory cells exiting GCs still follow the immunodominance hierarchy (Figure S2C-D). How does the immunodominance hierarchy become less pronounced in order to produce a balanced net of antibody titers directed toward different epitopes? The answer lies in the affinity-dependent expansion of memory B cells in the EGC, which has a more multi- clonal structure at all times (Figure 5D). At early times, most clones in the EGC target the immunodominant epitope. However, unlike in the GC, clones targeting subdominant epitopes that enter the EGC already have high affinity and can thus effectively compete with the immunodominant clones. Since cells in EGCs do not mutate, clones targeting subdominant epitopes remain competitive as time ensues. Thus, changes in antigen availability due to epitope masking give subdominant clones even more of a competitive advantage in EGCs than in GCs. As long as a few high affinity subdominant clones can enter the EGC and the EGC persists during therapy, they can expand to take over a significant portion of the EGC population. In contrast, many high affinity B cell clones targeting the immunodominant epitope continue to enter the EGC without a clear winner emerging. Thus, the clonal diversity of memory B cells targeting the immunodominant epitope is larger than those targeting subdominant epitopes (Figure 5E), which is a testable prediction.

### Time to rebound after ART cessation is determined by the characteristics of the aAb response

We next explored how aAbs that evolve during bnAb therapy could help confer ART-free viral control by modeling the virus dynamics post-ATI. Our goal is not to predict precise rebound times, but rather how the time to rebound depends on the characteristics of the generated aAb responses. Building upon prior models of HIV infection,^82–88^ we studied a “TIV” model where virions (V) infect target CD4+ T cells (T), are produced by infected cells (I), and mutate to escape antibody responses directed towards a number (*n*_*ep*_) of epitopes. Such a TIV model can be reduced to an effective dynamics of only the productively infected cells (STAR METHODS, Figure 6A).^35,89^ The dynamics of a cell, *I*_*i*_, infected by viral strain *i* is given by:

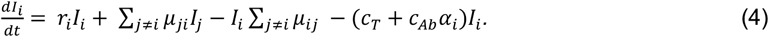

**Figure 6:**
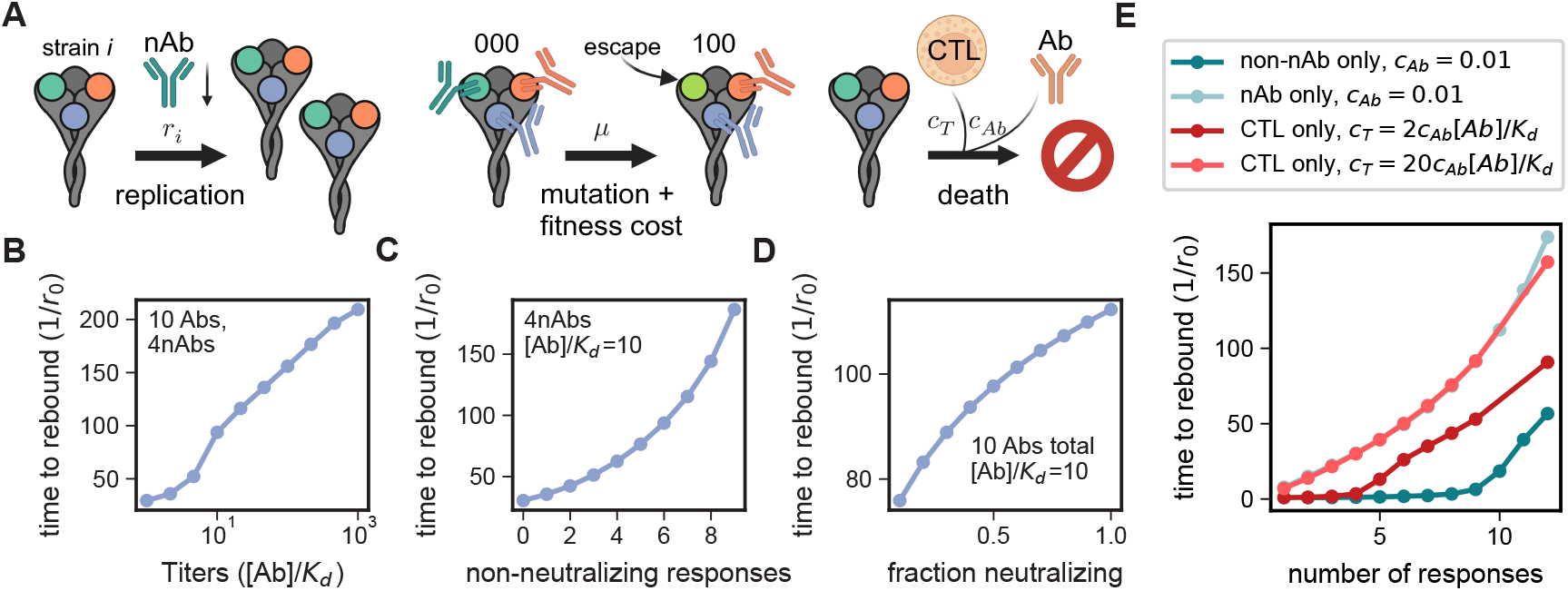
Viral dynamics after treatment interruption demonstrate that time to rebound depends on size and potency of antibody net targeting diverse epitopes. (A)Effective model of viral dynamics which takes into account the inhibition of replication/new infections by neutralizing aAbs and clearance of infected cells by CD8+ T cells / Cytotoxic T Lymphocytes (CTLs) and non-neutralizing effector functions of aAbs. Mutations can lead to escape from an antibody response toward a particular epitope with an associated fitness cost. (B-E) For the results shown, mutation rate *µ* = 0.005*r*_*0*_, the fitness cost for escaping any aAb response is *b*_*m*_ = 0.04, and *c*_*Ab*_ = 0.01*r*_*0*_. For (B-D), we chose *c*_*T*_ = 0.2*r*_*0*_. (B)Time to rebound increases with antibody titers targeting 10 distinct epitopes; aAbs targeting 4 epitopes are neutralizing. (C)Time to rebound increases with the number of responses directed toward non-neutralizing epitopes while keeping the number of neutralizing responses fixed at 4. (D)Time to rebound increases with the fraction of aAb responses that are neutralizing, with the total number of epitopes targeted fixed at 10. (E)Comparison of time to rebound if only non-neutralizing antibody responses (with titers [*Ab*]*/K*_*d*_ = 10 fo all epitopes), neutralizing antibody responses, or CTLs are in effect.

The effective rate at which new cells are infected, *r*_*i*_, is inhibited by neutralizing aAbs (nAbs) and the fitness cost, *b*_*m*_, due to evolving a mutation to evade aAb responses to epitope *m*. If we assume the total fitness cost is additive in the number of escaped epitopes, *r*_*i*_ is then given by (STAR Methods):

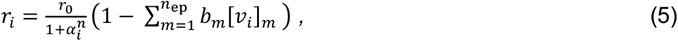

where 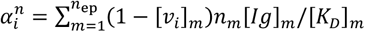 is the sum of nAb titers towards unescaped epitopes. Here, *n*_*m*_ = 1 if aAbs targeting epitope *m* are neutralizing and 0 otherwise, and [*v*_*i*_]_*m*_ = 1 if the strain has developed an escape mutation in epitope *m* and 0 otherwise. Mutations occur at a rate *µ*_*ij*_ = *µ* between pairs of viral strains separated by a mutation at one epitope. Infected cells are cleared by CD8+ T cells / Cytotoxic T Lymphocytes (CTLs) at a rate *c*_*T*_. Note that each viral population *i* in our model represents a collection of viral strains with the same antibody Env epitopes but potentially different T cell epitopes. Thus, *c*_*T*_ can be interpreted as a product of the clearance rate and the fraction of these strains with T cell epitopes that are targeted by CTLs during ATI. Infected cells are also cleared by antibody effector functions such as antibody-dependent cellular phagocytosis (ADCP) or antibody-dependent cellular cytoxicity (ADCC) at a rate *c*_*Ab*_*α*_*i*_, where 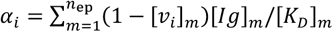 is the sum of antibody titers directed toward unescaped epitopes.^90,91^

Within our model, indefinite ART-free viral control is only possible if the fitness costs associated with escaping all aAb responses generated during therapy are sufficiently high, or if these strains can be controlled by CTL responses alone. However, many ART-free viral controllers following bnAb therapy did not show especially enhanced CD8+ T cell responses,^9^ and fitness costs for evolving mutations in Env epitopes are often not large.^35,92^ Therefore, we chose the parameters *b*_*m*_ and *c*_*T*_ to be in a regime where ART-free viral control would be lost if the viral swarm evolved mutations that escaped all the aAbs generated during therapy.

We initialize the population with 100 cells that are actively infected with the viral strain lacking any aAb escape mutations. We calculate the time to rebound as that required for the total population to reach twice its initial value.

We first consider a net of antibodies targeting 10 distinct epitopes, where the responses to 4 of these epitopes are neutralizing. For simplicity, we consider a case where fitness costs are the same for all epitopes, and antibody titers to each epitope are identical. As expected, we find that the time to rebound increases with antibody titers (Figure 6B). In addition, the time to rebound increases with the number of distinct epitopes targeted, even if the number of neutralizing epitopes targeted is held fixed (Figure 6C).

The increase is exponential because the number of paths from the initial state to the fully mutated state that evades all aAbs grows exponentially with the number of targeted epitopes. Since nAb responses can act via effector functions (ADCC, ADCP) and inhibit viral replication, we find that the time to rebound increases with the fraction of aAbs that target epitopes that result in neutralization (Figure 6D). Note that the results shown in Figure 6C-D are for antibody titers of 10, as data show that low titers of aAbs can be effective in blocking viral outgrowth.^35,38^

Next we tried to parse the relative roles of CTLs and neutralizing and non-neutralizing antibodies in ART-free viral control by examining the time to rebound post ART cessation if only one of these types of responses is in effect (Figure 6E). For a meaningful comparison of CTL responses and antibodies, we constructed a model where viral strains develop mutations to escape CTLs that target specific epitopes (analogous to Equation 4, but where strains are defined by their T cell epitopes instead of by their Env epitopes, see STAR Methods).

CTLs and non-neutralizing antibodies are similar in that they both act through effector functions. The relative rates of clearance, *c*_*T*_ versus *c*_*Ab*_, then determine which of these two responses are more effective at controlling viral load. In Figure 6E, we show two examples where *c*_*T*_ = 2*c*_*Ab*_[*Ig*]*/K*_*d*_ (2 times more potent than nonneutralizing antibodies) and *c*_*T*_ = 20*c*_*Ab*_[*Ig*]*/K*_*d*_ (20 times more potent than non-neutralizing antibodies). Our results show that the CTL clearance rate *c*_*T*_ must be significantly larger than the clearance rate due to antibody effector functions in order to delay viral rebound to the same extent as neutralizing antibodies (Fig. 6E, *c*_*T*_ = 20*c*_*Ab*_[*Ig*]*/K*_*d*_). This is because nAbs can act through effector functions and inhibit viral replication. This result suggests that neutralizing aAbs generated during therapy can contribute highly effectively to ART-free viral control post ATI.

## DISCUSSION

When administered at about the same time as ART^11^ or right after ART interruption^9^, HIV-specific bnAbs can lead to ART-free viral control in some people with HIV. Although the degree of control is variable and observed in only some participants, bnAbs show promise as a safe and effective intervention that could lead to sustained ART-free control. Understanding the mechanisms by which bnAb therapy engages host immune responses to facilitate this control is critical for optimizing this therapeutic strategy.

We propose that bnAbs can potently bind many circulating autologous viral strains which results in the display of diverse epitopes on FDCs. While viral replication is low during ART, aAbs targeting different epitopes on Env can sequentially evolve due to epitope masking (Figure 2). This mechanism is consistent with clinical data showing improved antibody responses to autologous and heterologous viruses in people with HIV 6 months after treatment with 3BNC117.^14^ In addition, the generation of diverse antibody responses upon sequential immunization with SARS-Cov-2 and influenza vaccines has also been observed clinically.^49,51,55,80^ In the HIV therapy context, such a polyclonal net of aAbs that evolves during bnAb therapy could delay viral rebound after ART interruption, since the virus would have to develop escape mutations in multiple epitopes in order to evade the antibody response (Figure 6).

Testing this mechanism would require detailed, temporal measurements of the antibody serum in participants with ART-free viral control both during and after bnAb therapy. Mapping the epitopes targeted by aAbs would be critical for determining whether aAbs targeting different epitopes evolve, as we predict in this study, and which conditions promote their evolution. Several technologies could enable these measurements, such as site-specific mutagenesis and other high throughput techniques.^35,93^

Many details of our mechanistic model could be refined in future simulation studies and with more experimental data. For example, Garg et al. present an additional mechanism for why passive immunization with bnAbs improves autologous antibody responses.^81,94^ They posit that because bnAbs bind antigen potently, the pulling force required for a GC B cell to internalize antigen off FDCs is greater, thus increasing the selection force towards high affinity B cells. Although their studies only included a single epitope, the effect of more stringent selection would likely delay emergence of subdominant responses in our multi-epitope model.

Similarly, we have neglected the antigen presentation and Tfh dynamics in the presence of bnAbs. Other simulation studies have shown that limited T cell and antigen availability can limit clonal diversity in GCs,^95^ albeit for a single epitope, and that antigen availability can be extended by internal recycling of immune complexes within FDCs.^96^ Our model assumes that GCs and EGCs persist for the duration of therapy due to long-term antigen availability.^57–59,97^ While six-month long GCs have been observed in response to slow delivery of HIV antigens,^98,99^ and GCs last at least three months after a booster shot of the COVID vaccines,^100^ it is unknown how long the humoral response lasts in response to passive bnAb immunization of chronically infected individuals. Measuring the distribution of diverse antigens on FDCs and the time that antigen is retained on FDCs post bnAB therapy would require imaging lymph nodes in animal models.^50,97,101^ Such experiments could help determine the dosing scheme and combination of bnAbs that optimizes antigen delivery and subsequent evolution of an antibody net during treatment.

In addition, we have neglected evolution of circulating and re-activated latent viral strains during bnAb therapy in response to selective pressure from bnAbs and coevolving aAbs.^14,35,46^ Temporal analysis of the viral sequences in latently infected cells and plasma during bnAb therapy will be important for determining how coevolution impacts the aAb response and its efficacy in ART-free viral control.^46,102^ As such data becomes available, our model could be extended to include these effects.

Several conditions need to be satisfied in order for a protective net of antibodies to evolve during bnAb therapy. As discussed in Figure 3, the net will not evolve if viremia is too low or too high at the time bnAbs are administered. Determining the “goldilocks” regime of antigen displayed on FDCs will be important for optimizing a therapeutic scheme that promotes the evolution of a net of aAbs. In addition, antibodies that develop during bnAb treatment will only be effective against re-activated latent viruses if (1) these antibodies are cross-reactive and/or (2) the viruses in the latent reservoir are not significantly different in sequence from viruses in circulation at the time of bnAb therapy. Our results show that cross-reactive antibodies can evolve if there are several epitopes that are shared across antigens displayed on FDCs (Figure 4). Low provirus diversity can be achieved if therapy is started soon following infection. Note also that a more narrowly focused net could achieve viral control if the provirus diversity is especially limited. These predictions can be tested in a clinical trial that measures viral load, plasma diversity, and provirus diversity prior to bnAb therapy and as a function time. Such measurements have been done in the ART context, and the data from those studies are consistent with our model predictions.^36^

We find that neutralizing autologous antibody responses are likely to be more effective than CTL responses at delaying viral rebound after ART cessation since nAbs both inhibit replication and carry out effector functions (Figure 6E). More generally, our study, which is consistent with clinical data,^14,51,55^ suggests that the humoral immune system has evolved mechanisms to naturally generate diverse responses. Therapeutic approaches that aim to enhance the evolution of diverse aAbs may therefore be easier to achieve than approaches that attempt to reverse the natural tendencies of the immune system, such as targeting CD8+ T cell epitopes that are not naturally immunodominant given a person’s HLA haplotype.^103–107^

Our model also has broad implications for passive immunization with therapeutic antibodies in other contexts.^108–110^ Indeed, other viruses such as influenza mutate over a much slower timescale within the population. Thus, vaccination with monoclonal antibodies that bind potently and broadly to a cocktail of viral strains could similarly enhance host memory B cell and Ab responses to unseen variants. We hope that rigorous testing of our proposed mechanism will unlock novel approaches to vaccination and cure for both acute and chronic viral infections.

## ACKNOWLEDGEMENTS

DK was supported by a National Science Foundation Graduate Research Fellowship under grant No. 2141064 and an internal MIT Fellowship. AKC and EW acknowledge support from NIH grant number 4R33AI161805-04. SRL and SGD are supported by the National Institutes of Health Delaney AIDS Research Enterprise (DARE) Collaboratory (UM1 AI164560-01). SRL is also supported by The National Health and Medical Research Council (NHMRC; grant number 2026490) and the mRNA Victoria Research Accelerator Fund.

## AUTHOR CONTRIBUTIONS

AKC, SRL, SD, DK, and EW conceived the project; AKC and DK designed the computational strategy; DK carried out the computational studies and carried out data analyses; EW assisted in developing the computer code; AKC and DK analyzed and interpreted the data with contributions from SRL and SD; AKC, DK wrote the first draft of the paper; AKC, SRL, SD, and DK edited and revised the paper.

## SUPPLEMENTAL INFORMATION

**Document S1. Table S1**

Table S1. Simulation parameters and their default values. Related to STAR Methods.

## SUPPLEMENTAL FIGURES

**Figure S1:**
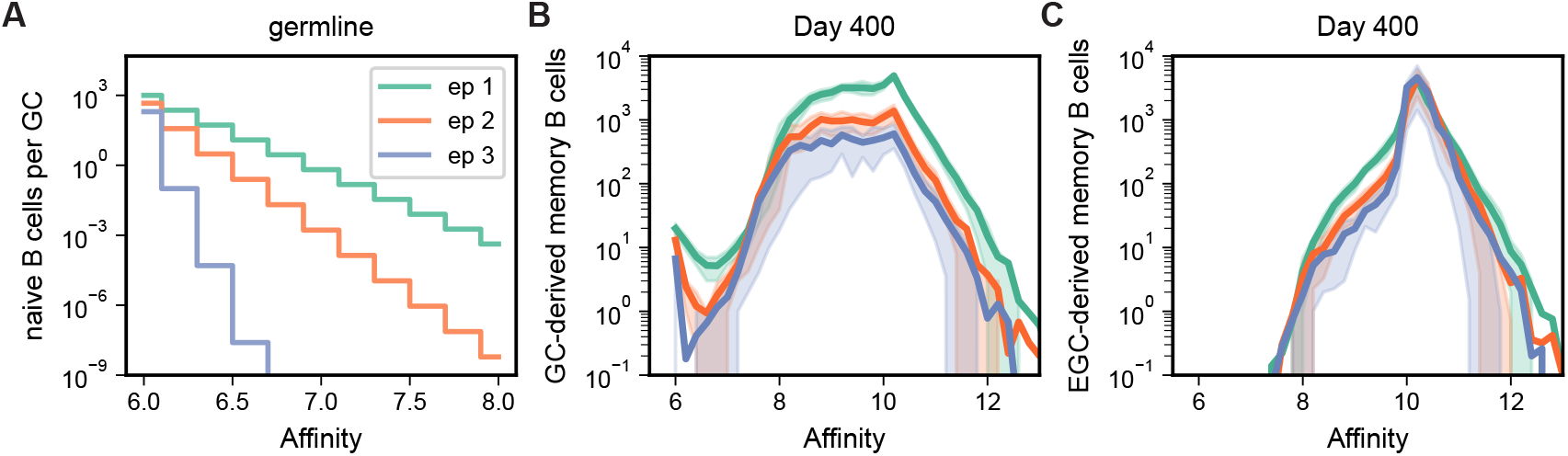
Affinity distributions of naive B cells and GC- or EGC-derived memory B cells. Binding affinity is reported in units of −log_01_(*K*_*d*_), where *K*_*d*_ is the dissociation constant. In panels B and C, thick lines show the mean over 50 simulation replicates and the shading is between the 25^th^ and 75^th^ percentiles. (A) Number of naive B cells in each affinity bin available to each GC at all times follows a truncated geometric distribution (Equation S2). (B) Affinity distribution of GC-derived memory B cells after 400 days of a sustained humoral immune response follows the immunodominance hierarchy. A small fraction of low-affinity memory cells are exported from the GC before they acquire sufficient affinity-enhancing mutations. (C) Affinity distribution of EGC-derived memory B cells after 400 days of a sustained humoral immune response is sharply peaked at the affinity ceiling of 10. Memory cells targeting all 3 epitopes have similar affinity distributions due to expansion of subdominant clones in the EGC.

**Figure S2:**
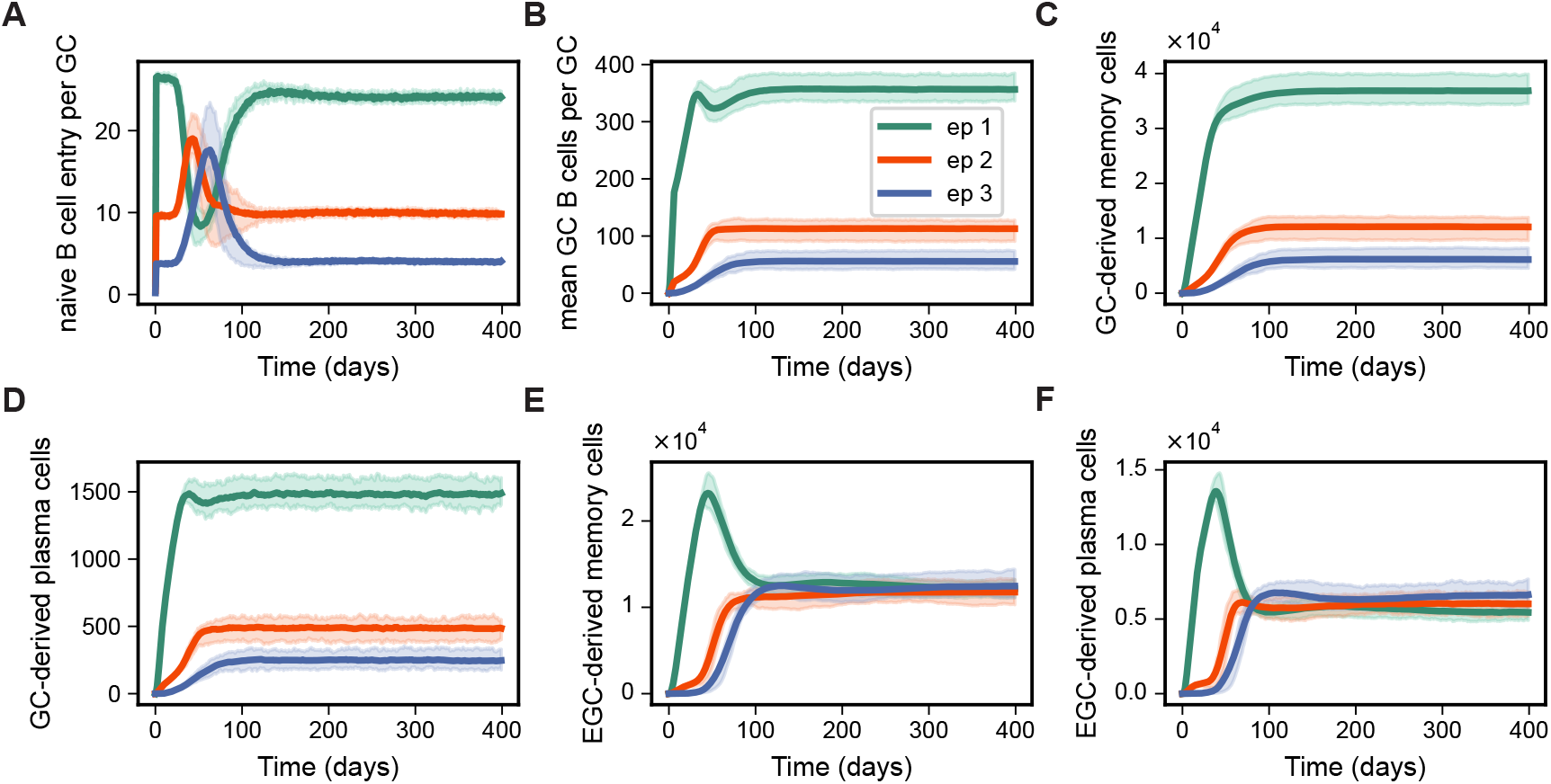
Dynamics of B cell populations during humoral immune response to bnAb therapy, related to Figure 2. Thick lines show the mean over 50 simulation replicates and the shading is between the 25^th^ and 75^th^ percentiles. All populations reach steady state within 200 days. (A) Mean number of naive B cells that enter a GC per day as a function of time (averaged over 200 GCs). (B) Mean number of GC B cells as a function of time (averaged over 200 GCs). (C)GC-derived memory cells and (D) GC-derived plasma cells follow the immunodominance hierarchy. (E) EGC-derived memory cells and (F) EGC-derived plasma cells targeting distinct epitopes reach similar population levels due to feedback regulation by autologous antibodies.

**Figure S3:**
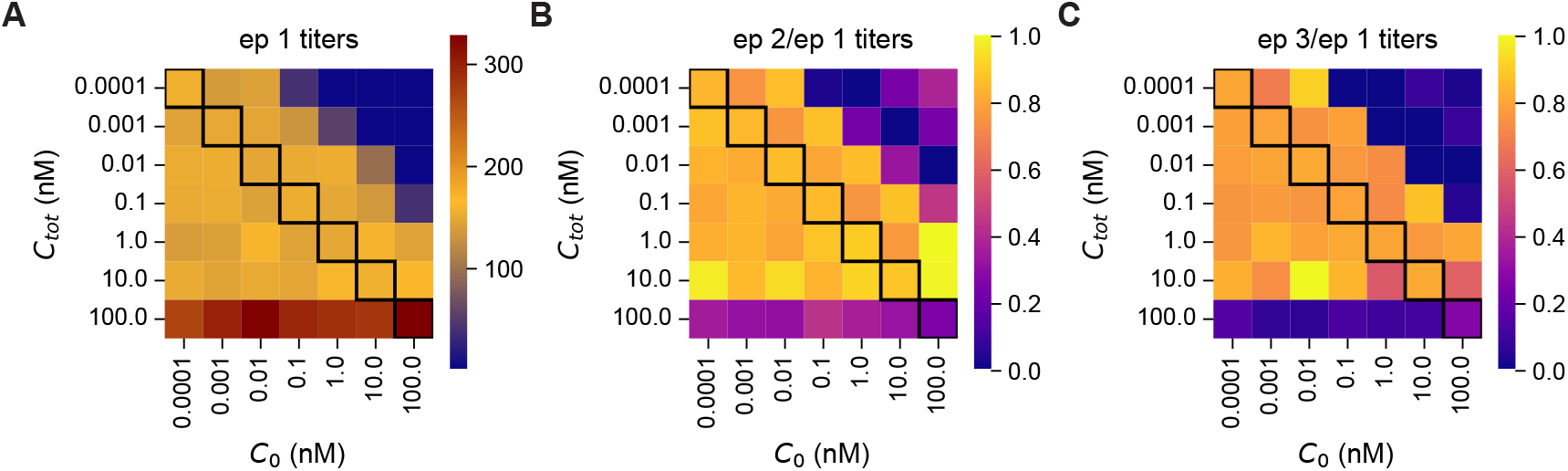
Optimal range of antigen concentration on FDCs for evolution of polyclonal net of aAbs targeting diverse epitopes depends on *C*_*0*_, related to Figure 3. (A) Titers ([Ab]/*K*_*d*_) of aAbs targeting the immunodominant epitope (ep 1) as a function of the total concentration of antigen on FDCs, *C*_*tot*_, and the reference concentration *C*_*0*_. Titers are low for small *C*_*tot*_/*C*_*0*_ due to lower activation probabilities of B cells at each selection step. (B) Ratio of titers towards epitope 2 and titers towards epitope 1 as a function of *C*_*tot*_ and *C*_*0*_. (C) Ratio of titers towards epitope 3 and titers towards epitope 1 as a function of *C*_*tot*_ and *C*_*0*_. Balanced titers towards all 3 epitopes occur at an intermediate range of antigen concentration on FDCs for a given *C*_*0*_. This range increases as *C*_*0*_ decreases since selection is less stringent.

**Figure S4:**
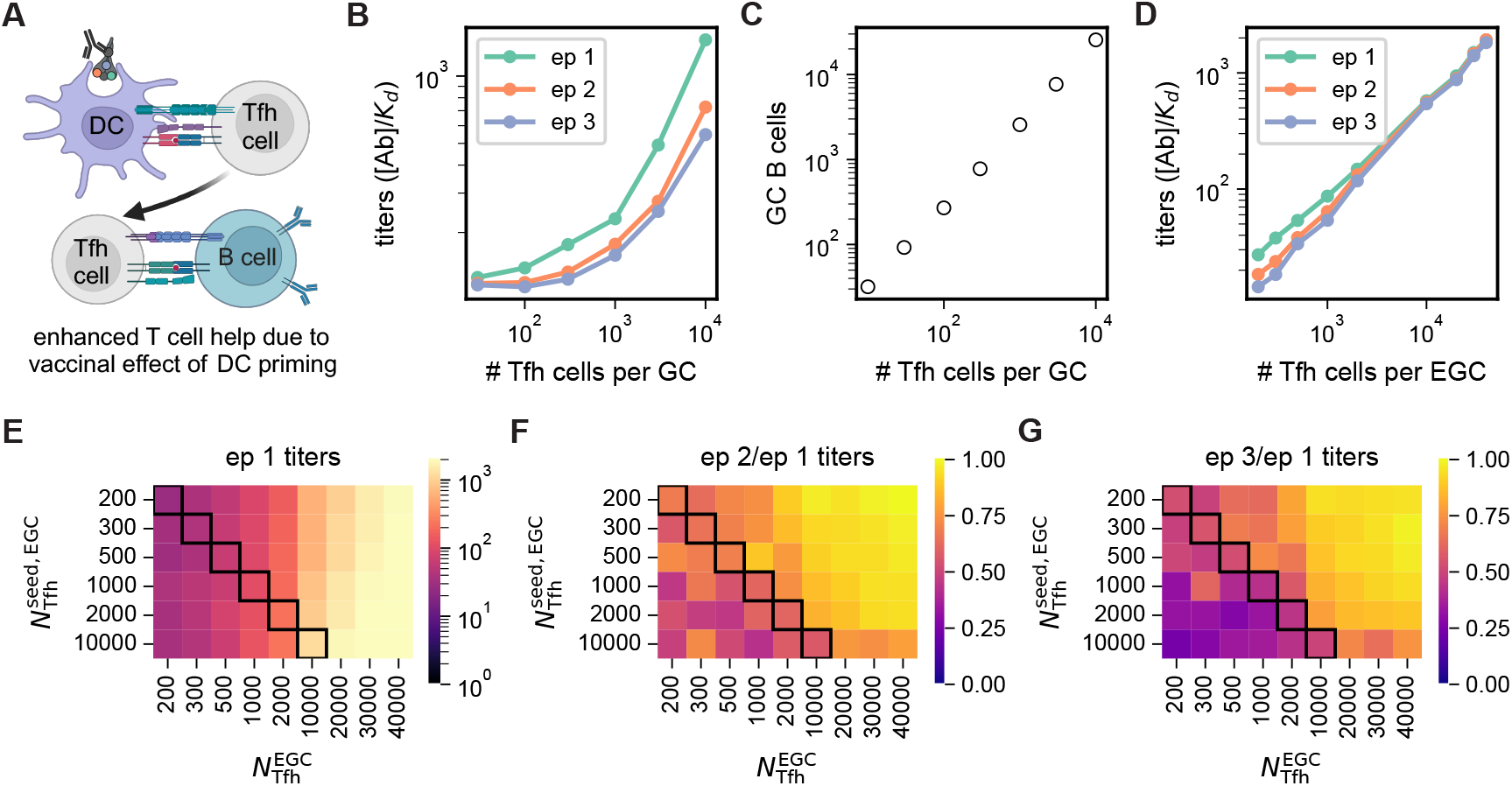
Effect of T follicular helper (Tfh) cells on antibody titers. (A) According to the “vaccinal” effect hypothesis, DCs are more effectively primed by bnAb-antigen immune complexes, resulting in enhanced activation of T cells, including Tfh cells. (B) Titers ([Ab]/*K*_*d*_) of aAbs targeting all 3 epitopes increase with the number of Tfh cells in each GC. (C) Number of GC B cells (averaged over 2000 GCs) increases linearly with the number of Tfh cells in each GC. (D) Titers of aAbs targeting all 3 epitopes increase with the number of Tfh cells in the EGC. (E) Titers of aAbs targeting the most immunodominant epitope (epitope 1) as a function of the number of Tfh cells involved in selection of memory B cells for EGC entry 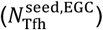 and the number of Tfh cells involved in selection of memory B cells for expansion within the EGC 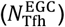. (F) Ratio of titers towards epitope 2 and titers towards epitope 1 as a function of 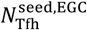 and 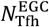. (G) Ratio of titers towards epitope 3 and titers towards epitope 1 as a function of 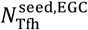 and 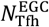. Titers across the 3 epitopes become more balanced and increase in scale as 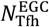 increases.

**Figure S5:**
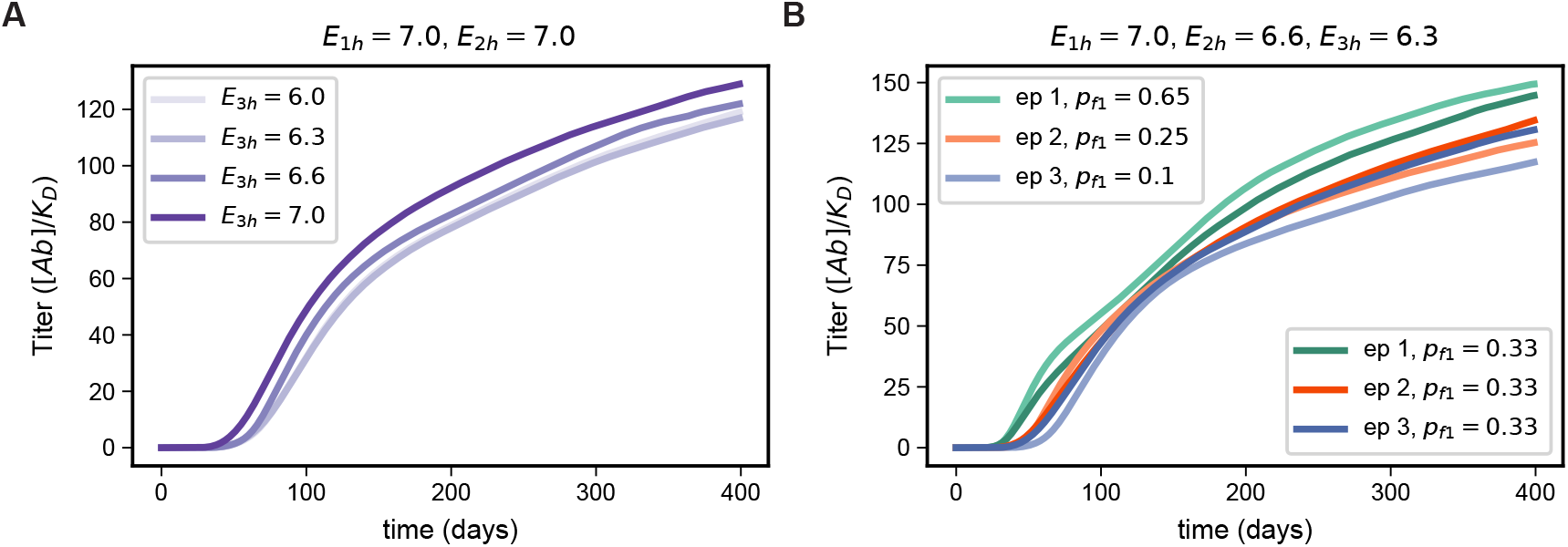
Effect of germline B cell characteristics on dynamics of subdominant responses. (A) Titers ([Ab]/*K*_*d*_) of aAbs targeting the least dominant epitope (ep 3) as a function of time for different values of *E*_3*h*_, with *E*_0*h*_ = *E*_2*h*_ = 7.0 fixed. Increasing the gap in germline affinity between germlines targeting epitope 3 and those targeting epitopes 1 or 2 delays the evolution of the most subdominant response. (B) The delay between responses is reduced when the fractions of naive B cells that target each epitope are more balanced (*p*_*f1*_ = *p*_*f*2_ = *p*_*f*3_ = 0.33, darker lines).

## STAR 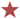 METHODS

### METHOD DETAILS

#### Simulations of the Humoral Immune Response

The computational model that we develop to study the humoral immune response in people living with HIV is adapted from our past work on the antibody response to sequential immunization with SARS-Cov-2 vaccines.^49^ Each simulation consists of 200 ongoing GC reactions and 1 EGC, which together define the dynamics of the following B cell populations: naive B cells, GC B cells, memory B cells, and antibody-secreting plasma cells. The memory and plasma cells are further divided into GC-derived populations and EGC-derived populations for bookkeeping purposes. Below, we describe the details of the simulation algorithm. The associated simulation parameters and their values are listed in Table S1.

##### Precursor frequencies and affinity distributions of germlineline B cell populations

###### Precursor frequencies

Each GC is associated with a pool of *N*_naive_ genetically distinct germline (or naive) B cells that are available to enter the GC. The value of *N*_naive_ can be determined from the total number of naive B cells in humans, which is estimated to be 10^01,111,112^ and the frequency of HIV-specific germline B cells, *p*_*f*_, as *N*_naive_ = *p*_*f*_ (1 × 10^01^)/(200 *GCs*). Of the *N*_naive_ B cells per GC, a fraction *p*_*f*1_ of them target epitope 1, a fraction *p*_*f*2_ of them target epitope 2, and so on, where 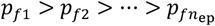.defines the hierarchy of precurosor frequencies, and 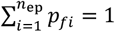.

For the three-epitope simulations, we assume an overall frequency of HIV-specific germlines of *p*_*f*_ = 1/(2.5× 10^4^).This number was chosen to be over an order of magnitude larger than the VRC01-specific precursory frequency, which is 1/(4 × 10^5^),^113^ and implies *N*_naive_ = 2000 cells per GC. We further chose *p*_*f1*_ = 0.65, *p*_*f*2_ = 0.25and *p*_*f*3_ = 0.1, which implies the number of germlines targeting the least dominant epitope (epitope 3) is 200, which is still larger than the number of VRC01-specific germlines, which is *N*_VRC10_ = 10^01^/(4 × 10^5^)/(200 GC*s*) = 125. This choice is motivated by the intuition that the number of precursors targeting a non-bnAb epitope should be larger than the number of precursors targeting the VRC01 epitope. We do not expect our qualitative results to change with different choices for the values of *p*_*f1*_ > *p*_*f*2_ > *p*_*f*3_. Results for balanced precursor frequencies across epitopes (*p*_*f1*_ = *p*_*f*2_ = *p*_*f*3_ = 1/3) can be found in Figure S5B.

For the simulations with 6 to 12 epitopes (Figure 4), we determined *p*_*f*_ in each case such that the total number of naive B cells targeting the least immunodominant epitope, 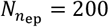.We also chose the number of precursors that target the most immunodominant epitope, *N*_1_, to be the same as in the 3-epitope case, i.e. 1300. We arranged the precursor frequencies for each epitope *p*_*f i*_ ∀*i* ∈ [1, *n*_ep_] in a geometric sequence 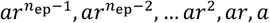.Here, *n*_ep_ is the total number of distinct epitopes, which can vary from 6 to 12 in the examples shown in Figure 4. We solve for the ratio between subsequent precursor target fractions, 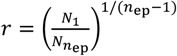 with *N*_1_= 1300 and 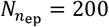 and then determine *a* using the constraint that 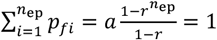. Once *a* and *r* are specified, the total number of naive B cells per GC is 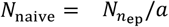. Thus, the number of naive B cells is different for each value of *n*_ep_.

###### Germline binding affinities

At the start of the simulation, each naive B cell is initialized with a germline binding affinity *E*_*j*_ = −log_01_(*K*_*d*_), where *K*_*d*_ is the dissociation constant and the affinities can take on one of 11 discrete values, *E*_*k*_ = 6 + δ*Ek* for δ*E* = 0.2 and *k* = 0, 1, …, 10. The minimum value *E*_0_ is chosen to match experimental observations of a minimum affinity required for naive B cell activation.^61,62^ The maximum affinity *E*_01_ = 8 is chosen based on the observation that naive B cell affinities can span two orders of magnitude.^114,115^ The probability of naive B cell *j* binding to epitope *i* with affinity *E*_*k*_, (*P*_*i*_(*E*_*j*_ = *E*_*k*_=)), is determined according to a geometric distribution for the germline B cell population targeting this epitope,

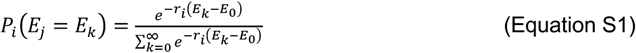

with slope *r*_*i*_. The expected number of naive B cells per GC in affinity bin *E*_*k*_ is this probability multipled by the total number of naive B cells that target a particular epitope,

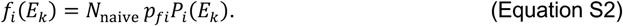

To determine the slope of the distribution, *r*_*i*_, we define a parameter, *E*_*ih*_, for each epitope-targeting germline population *i* such that *E*_*ih*_ corresponds to the affinity bin associated with probability *P*_*i*_(*E*_*ih*_) = 1/*N*_naive_. That is, it is unlikely to realize naive B cells with higher affinities than *E*_*ih*_. To further enforce the immunodominance hierarchy, this parameter is larger for more immunodominant epitopes, i.e. 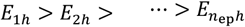.For the three-epitope simulations, we chose *E*_1*h*_ = 7, *E*_2*h*_ = 6.6, *E*_3*h*_ = 6.2. For the simulations with 6-12 epitopes, we chose these values to be linearly spaced between 7.0 and 6.0.

Using Equation S1 and *P*_*i*_(*E*_*ih*_) = 1/*N*_naive_, we obtain:

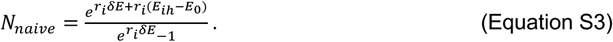

We can solve for *r*_*i*_ using Equation S3. Using this value of *r*_*i*_ and Equations S1 and S2 then gives the values of *f*_*i*_(*E*_*k*_). Since *f*_*i*_(*E*_*k*_) can lead to non-integer numbers of naive B cells, we add a uniform random number, *u* ∼ *U* (0, 1), and apply the floor function to obtain the final integer number of naïve B cells in each affinity bin,

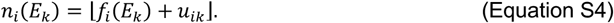

###### Mutation state vectors and binding affinity changes upon mutation

Each B cell *j* is assigned a mutation state vector, ***δ***_*j*_, of length *n*_res_= 80 indicating the presence (1) or absence (0) of a mutation at each residue of the B cell paratope. Naive B cells are initialized with vectors of all 0’s, indicating the germline state with affinity 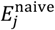.During an affinity-changing mutation step in the GC, a residue is selected at random to flip its value from 0 to 1 or 1 to 0. The affinity of the mutated B cell is then given by

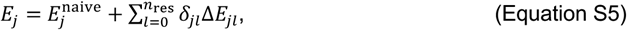

where Δ*E*_*jl*_ is the change in binding affinity associated with mutation in residue *l* in B cell *j*, and δ_*jl*_is the updated mutation state of residue *l* (1 or 0). For ease of computation, the values of Δ*E*_*jl*_ are precalculated at the start of the simulation. Based on previous work,^49,72^ we assume changes in the dimensionless Gibbs binding free energy are drawn from a shifted log-normal distribution,

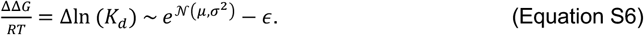

Since *E*_*j*_ = −log_01_(*K*_*d*_) is defined in base 10 units, this implies

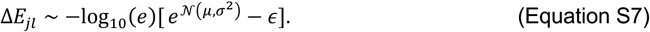

The parameters μ, *σ*^2^, and ϵ are chosen so that 5% of mutations are beneficial (Δ*E* > 0) and 95% of mutations are deleterious (Δ*E* < 0) (Table S1).

##### Antigen and antibody dynamics

###### Mapping from antigens to epitopes

We assume that antigen has already been deposited on FDCs by bnAbs prior to the start of the simulation. We further assume that the total antigen concentration on FDCs is fixed to some number, *C*_tot_, which is the sum of the concentrations of *n*_ag_different viral strains present on FDCs. We also assume that each antigen has some number of epitopes which can either be conserved between antigens or distinct. Distinct epitopes are associated with distinct germline populations, meaning a naive B cell that targets epitope 1 on antigen 1 will not be able to target epitope 1 on antigen 2 if it is different. Conserved epitopes have shared germlines. The total number of different epitopes present is then *n*_ep_ = *n*_ag_*n*_distinct_ + *n*_conserved_ where *n*_distinct_ and *n*_conserved_ are the number of different and shared epitopes on the antigens, respectively.

The concentration of each epitope on FDCs is specified by first defining a *n*_ep_ × *n*_ag_ matrix, ***R***, where *R*_*i*ℓ_ = 1 if epitope *i* is present on antigen ℓ and 0 otherwise. The total concentration of epitope *i* on FDCs is then

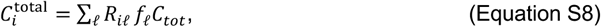

where *f*_ℓ_ is the fraction of antigen ℓ in circulation at the time of bnAb administration. For the three epitope simulations, we assume there is only 1 antigen with 3 epitopes. For the simulations associated with Figure 4, we consider the case of 2 antigens with 6 epitopes each present at equal concentrations on FDCs (*f*_0_ = *f*_2_ = 0.5), but vary the number of conserved epitopes from 0 to 6.

###### Antibody concentration and affinity dynamics

Autologous antibodies targeting epitope *i* are produced by plasma cells at a rate *k*_Ig_ and also decay at a rate *d*_Ig_,

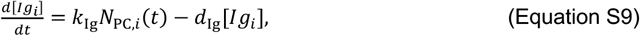

where *N*_PC,*i*_(*t*) is the number of GC and EGC-derived plasma cells specific to epitope *i* in circulation at a given time. Equation S9 is integrated forward in time with a first order Euler scheme.

The affinity *K*_*a,i*_ = 1/*K*_*d,i*_ of an antibody targeting epitope *i* is also updated at each time step. The governing equation can be derived using the product rule^49^

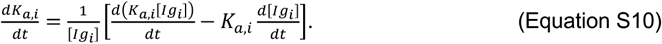

The first term within brackets above is the time derivative of the antibody titer, which is increased by new antibodies produced by plasma cells with mean affinity 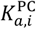,and decreased due to decay of free aAbs with affinity *K*_*a,i*_. If we also substitute Equation S9 into the second term, we get

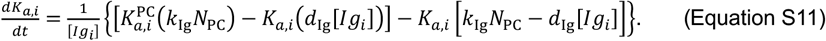

Thus, the time derivative of the affinity of an antibody targeting epitope *i* is given by

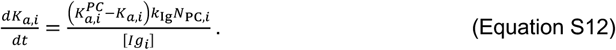

Note that when applying forward Euler to Equation S12, we use the approximation

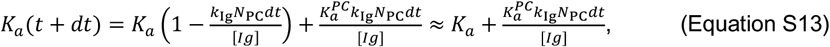

since *k*_Ig_*N*_PC_*dt ≪* [*Ig*].

###### Epitope Masking

The effective concentration of each epitope presented on FDCs that is available to B cells changes over time due to epitope masking by autologous antibodies. To determine the concentration, 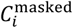,of the immune complex formed by an autologous antibody with effective concentration [*Ig*_*i*_]^∗^ and its target epitope *i*, we consider the following binding equilibrium:

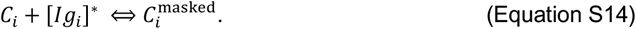

Since 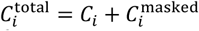,which is conserved throughout the simulation, and defining 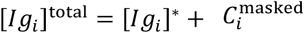, we obtained the masked epitope concentration by solving the following quadratic equation

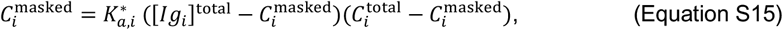

where 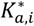 is the effective binding affinity of the antibody to its target epitope *i*. In turn, the concentration of unmasked antigen which is available to B cells on FDCs is

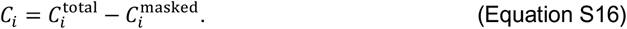

If the epitopes are spatially non-overlapping, then [*Ig*_*i*_]^∗^ is simply the free concentration of aAbs targeting epitope *i*. However, in the presence of epitope overlap, other antibodies can also partially bind to epitope *i*. We take this into account by defining (in the 3-epitope case)

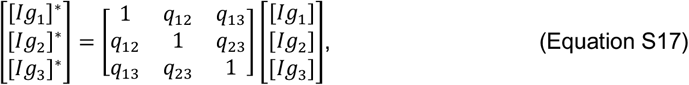

where *q*_*mn*_ is the amount of overlap between epitopes *m* and *n*. Similarly, we also adjust the effective binding affinities to account for epitope overlap

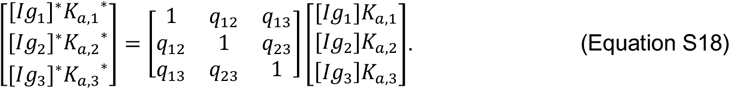

Detailed structural data on a system would allow more precise determination of the quantities calculated using Equations S17 and S18.

##### GC & EGC Dynamics

###### Simulation Algorithm

After all germline B cell populations are initialized at the start of the simulation, the simulation of the humoral immune response begins. At the beginning of each time step, the unmsked concentration, *C*_*i*_, of each epitope on FDCs is calculated using Equation S16. Then, naive B cells from each GC pool are stochastically activated with probability *P*_ativated_ (Equation 1) which depends on the amount of antigen internalized, which in turn depends on *C*_*i*_ and the germline binding affinity *E*_*j*_ (Equation 2). Activated naive B cells then compete for productive interactions with Tfh cells and are selected with probability *β*_*j*_*dt*, where *β*_*j*_ is the selection rate (Equation 3). Activated and selected B cells then enter each of the 200 independent GCs. Note that all *N*_naive_ lineages of naive B cells associated with each GC are always available for activation in every time step, i.e. we assume the naive B cell pool does not deplete over the course of the simulation. Upon entering a GC, each B cell divides twice so that 4 identical B cells are added to the GC.

In each time step, GC B cells undergo one round of selection, proliferation, export, and mutation. First, GC B cells are positively selected in two steps (antigen internalization and selection by T cells, Equations 1-3). Positively selected B cells give birth, resulting in two copies of daughter cells. One copy does not mutate or leave the GC. Daughter cells in the second copy exit the GC with probability 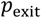.The exported cells have a probability 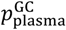 of differentiating into GC-derived plasma cells and a probability 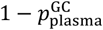 of differentiating into GC-derived memory cells. After day 6, all the non-exported daughter cells in the second copy mutate (but since the first copy does not, effectively, daughter cells have a probability of 0.5 of mutating).^72^ This is an overestimate of the mutation rate. Prior to day 6, GC B cells do not mutate. Mutations have a probability of being silent, *p*_silent_, or lethal, *p*_lethal_. Nonsilent, nonlethal mutations lead to a change in the binding affinity of the GC B cell, according to Equations S5 and S7.

In each time step, GC-derived memory B cells are selected for entry into the EGC in two steps (Equations 1-3). In the context of this work, we do not allow for the possibility that GC-derived memory cells could re-enter the GC. In past work, we have seen that allowing some fraction of memory B cells to re-enter GCs attenuates sub-dominant responses.^49^ In each time step, EGC B cells undergo one round of selection, proliferation, and export. First, EGC memory B cells are positively selected in two steps (Equations 1-3). Then, positively selected EGC memory B cells give birth, where one daughter cell remains in the EGC and the other is exported 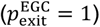.The exported daughter cells have probability 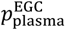 of differentiating into an EGC-derived plasma cell and probability 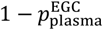 of being re-added to the memory EGC B cell population.

After one round of each of the 200 GC reactions and 1 EGC reaction, all B cell populations (GC B cells, and GC- or EGC-derived plasma and memory B cells) undergo apoptosis at different rates based on experimental measurements of B cell half lives (see Table S1). We can convert a B cell half life *t*_0/2_ to a death rate using

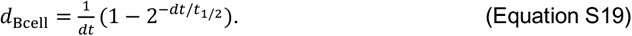

At the end of the simulation timestep, the concentrations of the antibodies and and their affinities to their target epitopes are updated using Equations S9 and S12. All of the above computational steps are repeated in the next time step.

One simulation corresponds to 400 days of a sustained humoral immune response with 200 ongoing GCs and 1 ongoing EGC (each of which have continuous entry and exit). Thus, all B cell populations reach a steady state determined by a balance of immigration, emigration, birth, and death rates (Figure S2). Below, we describe how various parameters were chosen to achieve reasonable steady state population levels and antibody titers.

###### Selection-associated Parameters

There are 4 selection steps that occur in the simulation: selection of naive B cells for GC entry [seed, GC], selection of GC B cells within a GC [GC], selection of GC-derived memory cells for EGC entry [seed, EGC], and selection of memory EGC B cells within the EGC [*E*GC]. All 4 selection steps involve activation by antigen internalization (Equations 1-2), which depends on the maximum positive selection rate *β*_0_, and competition for T cell help (Equation 3), which depends on the ratio of the number of Tfh cells, *N*_Tfh_, and the number of B cells in the population at a given time. Thus, each selection step has 2 associated parameters. In principle, the Tfh cells may also have dynamics, but in the absence of experimental measurements, we assume the amount of Tfh associated with each selection step is constant.

The first selection step controls the rate of entry of naive B cells into the GC. Although this quantity has not been measured, we chose the selection rate 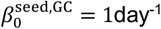 and 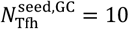 so that ∼30-50 B cells enter the GC per day (Fig. S2A). Increasing 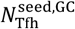 too much floods the GC with naive B cells and hinders affinity maturation.

For the second selection step inside GCs, we chose 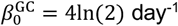 based on the observation that the average GC B cell divides 4 times in a day (cell cycle of 6 hours).^69^ Note that the size of the steady state GC B cell population is primarily determined by a balance of the GC B cell birth rate, death rate, and 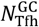.Observations from fine needle biopsy (FNB) measurements of lymph nodes in people living with HIV suggest that the average ratio of GC B cells to Tfh is roughly 2.5 and that the average number of GC B cells is ∼10^5^ in each FNB sample in ART-naïve people (which implies ∼500 B cells per GC).^70^ We thus chose 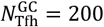 in each GC and adjusted the GC B cell death rate *d*_GC_ to ensure a population of ∼500 B cells per GC.

For selection of GC-derived memory B cells for EGC entry, we chose 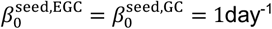 and varied 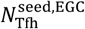 to study the effect of this parameter on the antibody titers. These parameters control the rate at which memory cells enter the EGC, 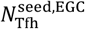 does not have a significant effect on the scale of antibody titers (Figure S4E). Increasing 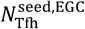 makes the antibody titers less balanced across epitopes (Figure S4F-G) since it floods the EGC with GC-derived memory B cells, which follow the immunodominance hierarchy (Figure S2C). The main text results use 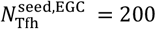.

The maximum positive selection rate of EGC memory B cells is assumed to be the same as that of a GC B cell, 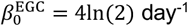.Increasing the number of T cells in the EGC, 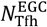,increases the overall scale of the antibody titers (Figure S4D-E) and also promotes more balanced titers across epitopes (Figure S4D, F-G). We chose 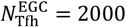 to get antibody titers on the order of 100, consistent with antibody titer measurements from participants in the VISCONTI study.^45^

#### Viral Dynamics Model

Building upon prior models of HIV infection, we studied a “TIV” model of viral dynamics in response to immune pressure.^82,83^ Each viral strain *i* is specified by a mutation state vector ***v***_***i***_ of length *n*_*ep*_ where [*v*_*i*_]_*m*_ = 1 if the strain develops an escape mutation in Env epitope *m*, and equals zero otherwise.^84,85^ Virions of each strain, *V*_*i*_, infect target CD4+ T cells at a rate *β*_*i*_, are produced by infected cells (*I*_*i*_) at a rate *p*_*i*_, are cleared at a rate δ, and mutate from strain *i* to strain *j* at a rate μ_*ij*_. Additionally, infected cells are cleared by CTLs at a rate *c*_*T*_. Assuming the reservoir of target cells is constant, *T*, at the start of ATI, and that mutation is uncoupled from replication, one arrives at the following system of ODEs describing the dynamics of each of the 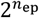 populations of viruses and infected cells if there are no antibody responses,

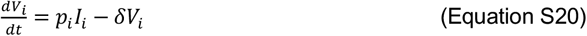

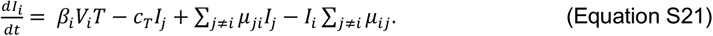

Antibodies targeting epitope *m* (*Ig*_*m*_) bind to free virions of strain *i* with affinity 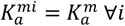 to form immune complexes, *Ig*_*m*_ + *V*_*i*_ → *Ig*_*m*_ − *V*_*i*_, as long as the viral strain, *i*, has not developed an escape mutation in epitope *m*; i.e., [*v*_*i*_]_*m*_ = 0. We assume immune complex formation is fast, resulting in 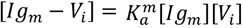. In our model, both neutralizing and non-neutralizing antibodies can bind to free virions. Defining 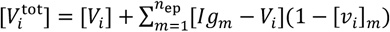, one can show that the concentration of unbound virions is given by

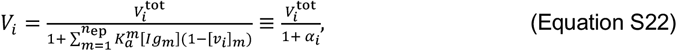

mirroring the functional form in Refs.^86,87^. The non-neutralizing effector functions (ADCC and ADCP) of neutralizing and non-neutralizing antibodies are taken to act on infected cells (e.g., by binding to budding virions) at a rate proportional to their binding affinity to their target epitope, *c*_Ab_*α*_*i*_.^90,91^

Antibodies targeting neutralizing epitopes are defined by an indicator function, *n*_*m*_ = 1, which is 0 otherwise. We mimic the ability of neutralizing antibodies to prevent infection by enhancing the viral clearance rate δ by an amount proportional to the total antibody titers directed towards unmutated neutralizing epitopes,^81,87^

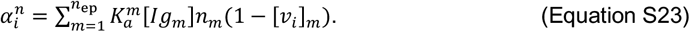

Incorporating clearance of free virus by neutralizing antibodies, substituting Equation S22 into S20, and assuming rapid formation of immune complexes gives the dynamics of the total concentration of viral strain *i*,

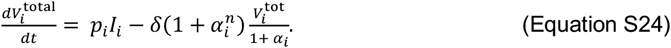

Due to the rapid clearance of virus particles from circulation, viral dynamics are much faster than that of cells.^116^ We can thus apply the adiabatic approximation, 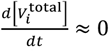 to Equation S24, which gives

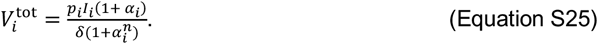

Substituting Equations S25 and S22 into Equation S21 gives the effective dynamics of infected cells in Equation 4:

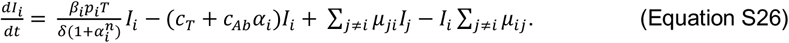

Note that the effective replication rate of infected cells in Equation S26 could equivalently be derived by reducing the infection rate *β*_*i*_ by a factor 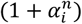.^88^ We further define a baseline rate *r*_1_ = *β*_1_*p*_1_*T*/δ at which infected cells are infected with the wildtype strain with no escape mutations ([*v*_1_]_*m*_ = 0 ∀*m*).^35^ All other strains incur fitness costs associated with each escape mutation, *b*_*m*_, which we assume are additive in the number of escape mutations.^89^ Thus, the effective replication rate of infected cells is given by

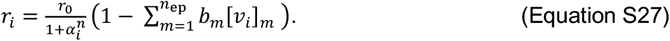

##### Comparison model of viral dynamics in response to CTL pressure alone

For the purposes of comparing CTLs to antibodies, we also construct an alternative model where viral strains are instead defined by a mutation state vector ***w***_*i*_ where [*w*_*i*_]_*m*_ = 1 if the virus develops an escape mutation in T cell epitope *m*. We again assume that an escape mutation in a T cell epitope incurs a fitness cost *b*_*m*_. If all CTLs clear infected cells at the same rate *c*_*T*_ then the effective dynamics of infected cells are

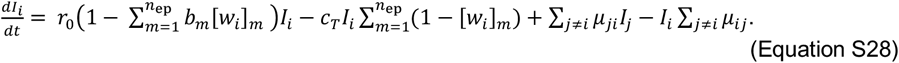

Equation S28 was used to calculate the time to rebound as a function of the number of CTL responses (*n*_ep_) in Fig. 6E.

