## Supplementary material for "Mechanism for evolution of diverse autologous antibodies upon broadly neutralizing antibody therapy of people with HIV": Table S1

**Table S1: Simulation parameters and their default values. Related to STAR Methods.**

| Parameter | Default Value | Description | Note |
|---|---|---|---|
| **Antigen and antibody dynamics** | | | |
| $dt$ | 0.05 day | Time step for simulations | Same as in Ref.[1] |
| $k_{\mathrm{Ig}}$ | $2.5 \times 10^{-5}$ nM day$^{-1}$ PC$^{-1}$ | Rate of antibody production per plasma cell (PC) per day. | Based on measured mean secretion rate of 174 IgG s$^{-1}$ PC$^{-1}$ and mouse TBV of ~1 ml.[2] |
| $d_{\mathrm{Ig}}$ | 0.01 day$^{-1}$ | Antibody decay rate | Based on antibody half life of ~28 days[3], $d_{\mathrm{Ig}} = \log(2)/28$ |
| $[Ig_i]_0$ | 0.01 nM | Initial aAb concentration for each epitope $i$ | Same as in Ref.[1] |
| $K_{a,i}$ | 0.001 nM$^{-1}$ | Initial aAb affinity for each epitope $i$ | Same as in Ref.[1] |
| $C_{\mathrm{tot}}$ | 1.0 nM | Fixed total antigen concentration on FDCs | Varied in Fig. 3 and Fig. S3 |
| $n_{\mathrm{ag}}$ | 1 | Number of antigens on FDCs | Varied to $n_{ag} = 2$ in Fig. 4 |
| $f_\ell$ | 1 | Fraction of antigen $\ell$ on FDCs | Varied to $f_1 = f_2 = 0.5$ in Fig. 4 |
| $n_{\mathrm{conserved}}$ | 0 | Conserved epitopes across antigens | Varied from 0 to 6 in Fig. 4 |
| $n_{\mathrm{distinct}}$ | 3 | Distinct epitopes across antigens | Varied from 0 to 6 in Fig. 4 |
| masking | 1 | Whether to turn on epitope masking | Varied (1 or 0) in Fig. 2D |
| **Parameters specific to three-epitope simulations** | | | |
| $N_{\mathrm{naive}}$ | 2000 | Number of naïve B cells per GC | ~$10^{10}$ total naive B cells[4,5] times estimated HIV-specific precursor frequency of ~1 in $2.5 \times 10^4$ divided by 200 GCs |
| $p_{f1}$ | 0.65 | Fraction of naïve B cells that target epitope 1 | Varied to $p_{f1} = 0.33$ in Fig. 5C and Fig. S5B |
| $p_{f2}$ | 0.25 | Fraction of naïve B cells that target epitope 2 | Varied to $p_{f1} = 0.33$ in Fig. 5C and Fig. S5B |
| $p_{f3}$ | 0.10 | Fraction of naïve B cells that target epitope 3 | Varied to $p_{f1} = 0.33$ in Fig. 5C and Fig. S5B |
| $E_1^h$ | 7.0 | Parameter for the germline affinity distribution of epitope 1 | Same as in Ref.[1] |
| $E_2^h$ | 6.6 | Parameter for the germline affinity distribution of epitope 2 | Set to $E_2^h = 7.0$ in Fig. S5A |
| $E_3^h$ | 6.2 | Parameter for the germline affinity distribution of epitope 3 | Varied in Fig. S5A |
| $q_{12}$ | 0 | Amount of spatial overlap between epitope 1 and 2 | Varied from 0 – 0.8 in Fig. 2F |
| $q_{13}$ | 0 | Amount of spatial overlap between epitope 1 and 2 | Varied from 0 – 0.8 in Fig. 2F |
| $q_{23}$ | 0 | Amount of spatial overlap between epitope 2 and 3 | |
| **GC and EGC dynamics** | | | |
| $C_0$ | 1.0 nM | Reference antigen concentration | Varied in Fig. S3 |
| $E_0$ | 6 | Minimum affinity for naïve B cell activation | Based on measurements of germline $K_d$ ≈1$\mu$M[6,7] of activated naive B cells |

| | | | |
|---|---|---|---|
| $E_{\mathrm{sat}}$ | 10 | Affinity ceiling at which amount of antigen internalized saturates[8] | Same as in Ref.[1] |
| $K$ | 0.5 | Stringency of selection by Tfh cells | Same as in Ref.[1] |
| $n_{\mathrm{res}}$ | 80 | Length of mutation state vector | Based on upper range of sum of CDR lengths in light and heavy chain [9] |
| $p_{\mathrm{lethal}}$ | 0.3 | Probability that a mutation is lethal[9] | Same as in Ref.[1] |
| $p_{\mathrm{silent}}$ | 0.5 | Probability that a mutation is silent[9] | Same as in Ref.[1] |
| $\mu$, $\sigma$, $\epsilon$ | 3.1, 1.2, 3.08 | Parameters for shifted log-normal distribution that models the effect of affinity-changing mutations | Same as in Ref.[1] |
| $N_{\mathrm{Tfh}}^{\mathrm{seed,GC}}$ | 10 | Tfh cells involved in selection of activated naïve B cells for GC entry | Chosen so that 30-50 naive B cells enter each GC per day |
| $N_{\mathrm{Tfh}}^{\mathrm{GC}}$ | 200 | Tfh cells involved in selection of activated GC B cells within GCs | Varied in Fig. S4 |
| $N_{\mathrm{Tfh}}^{\mathrm{seed,EGC}}$ | 200 | Tfh cells involved in selection of activated GC-derived memory B cells for EGC entry | Varied in Fig. S4 |
| $N_{\mathrm{Tfh}}^{\mathrm{EGC}}$ | 2000 | Tfh cells involved in selection of activated memory EGC B cells within EGC | Varied in Fig. S4 |
| $\beta_0^{\mathrm{seed,GC}}$ | 1 day$^{-1}$ | Maximum rate of selection for GC entry | Same as in Ref.[1] |
| $\beta_0^{\mathrm{GC}}$ | 2.77 day$^{-1}$ | Maximum rate of selection for GC B cells | Based on observation that GC B cells divide 4 times per day[10], $\beta_0^{GC} = 4\ln(2)$. |
| $\beta_0^{\mathrm{seed,EGC}}$ | 1 day$^{-1}$ | Maximum rate of selection for EGC entry | Same as in Ref.[1] |
| $\beta_0^{\mathrm{EGC}}$ | 2.77 day$^{-1}$ | Maximum rate of selection for EGC B cells | Same as in Ref.[1] |
| $d_{\mathrm{GC}}$ | 0.4 day$^{-1}$ | Death rate of GC B cells | Chosen so that average number of B cells in each GC is 500, for a total of $10^5$ GC B cells across 200 GCs, matching measurements from fine needle biopsies of lymph nodes in ART-naive people with HIV[11] |
| **Plasma and memory cell dynamics** | | | |
| $p_{\mathrm{exit}}^{\mathrm{GC}}$ | 0.05 | Probability that one of the daughter cells of a positively selected GC B cell exits and differentiates[12] | Same as in Ref.[1] |
| $p_{\mathrm{exit}}^{\mathrm{EGC}}$ | 1.0 | Probability that one of the daughter cells of a positively selected EGC memory B cell exits and differentiates | Same as in Ref.[1] |
| $p_{\mathrm{plasma}}^{\mathrm{GC}}$ | 0.1 | Probability that an exported GC B cell differentiates into a plasma cell (as opposed to memory) | Same as in Ref.[1] |
| $p_{\mathrm{plasma}}^{\mathrm{EGC}}$ | 0.6 | Probability that an exported EGC B cell differentiates into a plasma cell (as opposed to re-entering EGC) | Based on observation that 60% of reactivated memory B cells differentiate into plasma cells[13] |
| $d_{\mathrm{PC}}$ | 0.17 day$^{-1}$ | Death rate of plasma cells | Based on half life of 4 days[14] |
| $d_{\mathrm{mem}}$ | 0.06 day$^{-1}$ | Death rate of memory cells | Based on half life of 11 days[15] |
